# The CRISPR-Cas9 crATIC HeLa transcriptome: Characterization of a novel cellular model of ATIC deficiency and ZMP accumulation

**DOI:** 10.1101/2020.04.28.067066

**Authors:** Randall C Mazzarino, Veronika Baresova, Marie Zikánová, Nathan Duval, Terry G. Wilkinson, David Patterson, Guido N. Vacano

**Author notes:** Corresponding author: Guido Vacano, Ph.D., Address: University of Denver, 2155 E. Wesley Avenue Room 543, Denver, CO 80210.

## Abstract

In *de novo* purine biosynthesis (DNPS), 5-aminoimidazole-4-carboxamide ribonucleotide formyltransferase (2.1.2.3) / inosine monophosphate cyclohydrolase (3.5.4.10) (ATIC) catalyzes the last two reactions of the pathway: conversion of 5-aminoimidazole-4-carboxamide ribonucleotide [aka Z-nucleotide monophosphate (ZMP)] to 5-formamido-4-imidazolecarboxamide ribonucleotide (FAICAR) then to inosine monophosphate (IMP). ZMP is an adenosine monophosphate (AMP) mimetic and a known activator of AMP-activated protein kinase (AMPK). Recently, a HeLa cell line null mutant for ATIC was constructed via CRISPR-Cas9 mutagenesis. This mutant, crATIC, accumulates ZMP during purine starvation. Given that the mutant can accumulate ZMP in the absence of treatment with exogenous compounds, crATIC is likely an important cellular model of DNPS inactivation and ZMP accumulation. In the current study, we characterize the crATIC transcriptome versus the HeLa transcriptome in purine-supplemented and purine-depleted growth conditions. We report and discuss transcriptome changes with particular relevance to Alzheimer’s disease and in genes relevant to lipid and fatty acid synthesis, neurodevelopment, embryogenesis, cell cycle maintenance and progression, extracellular matrix, immune function, TGFβ and other cellular processes.

## 1. Introduction

*De novo* purine synthesis (DNPS) is essential for cellular function. Purines are critical as 1) components of DNA and RNA: the information carrying molecules of cells, 2) intra and intercellular signaling molecules, for example G (guanine) protein coupled receptors, 3) as the major source of energy currency in the form of ATP and 4) substrates and co-enzymes for many cellular functions. In humans and other animals DNPS is accomplished via ten sequential enzymatic reactions resulting in conversion of phosphoribosyl pyrophosphate (PRPP) to inosine monophosphate (IMP) mediated by six enzymes (seven of the ten reactions are catalyzed by three multifunctional enzymes). IMP can then be converted through two reactions to either AMP or GMP.

AMP-activated protein kinase (AMPK) is a major regulator of cellular metabolism. AMPK regulates the mammalian Target of Rapamycin (mTOR) pathway. AMPK activation shuts down mTOR signaling directly via phosphorylation of Raptor [1] and indirectly via activation and phosphorylation of Tuberous Sclerosis Complex 1 and 2 (TSC1 and TSC2). mTOR regulates vital cellular anabolic and catabolic processes involved in lipid synthesis, glycolysis, mitochondrial and lysosomal biosynthesis, apoptosis, glucose metabolism, cytoskeletal rearrangement, and cell migration [1]. Recently, AMPK activation has been implicated in different cellular processes such as inflammation suppression [2], and suppression of several IFN-γ-induced cytokines and chemokines in primary astrocytes and microglia via its restriction of IFN-γ signaling [3].

AMP is an allosteric effector of AMPK, and AMPK responds to increases in the AMP:ATP ratio by inhibiting ATP catabolism and promoting ATP anabolism [4,5]. AMPK can activate these processes in response to ATP depletion produced via fasting, exercise [6] or other means. Generally speaking, AMP has three roles in AMPK control: promotion of AMPK phosphorylation [7], inhibition of AMPK dephosphorylation [8], and allosteric activation of phosphorylated AMPK [9].

ZMP is an AMP mimetic and a potent AMPK agonist. While AMPK has garnered the most attention as the best characterized effector of ZMP accumulation, several studies have shown that ZMP has multiple cellular targets [7,10,11]. This supports the hypothesis that ZMP can allosterically regulate enzymes ordinarily regulated by AMP. In multiple studies analyzing ZMP function, cells were fed AICA riboside (AICAr, the dephosphorylated form of ZMP) which is converted to ZMP within the cell. This approach involves adenosine transporters and adenosine kinases to ensure ZMP production and accumulation in the cell [4], however, once ZMP is formed, cells with active ATIC enzyme will catalyze it via DNPS. Recent work indicates that AICAr itself affects cell metabolism independently of its role as substrate for ZMP synthesis. For example, AICAr administration induces a potentially deleterious intracellular ATP depletion in hepatocytes [12]. Drug intervention inhibiting ATIC dimer formation has been shown to be efficacious in reducing tumor cell proliferation [13], but can produce off target effects. Therefore, cell models that accumulate ZMP without exogenous AICAr should be advantageous for investigating optimal ZMP dosage levels in cells to produce beneficial effects.

The bifunctional enzyme 5-aminoimidazole-4-carboxamide ribonucleotide formyltransferase (2.1.2.3) / inosine monophosphate cyclohydrolase (3.5.4.10) (ATIC) catalyzes the final two reactions of IMP synthesis, converting ZMP to FAICAR and then FAICAR to IMP (Figure 1). It is a homodimer, and dimerization is required for activity [14]. Under some conditions ATIC may be a rate limiting step of the DNPS pathway [15,16]. So far, a single human with ATIC deficiency (AICA-ribosiduria [15]) has been identified and the mutations characterized: a mis-sense mutation in one allele resulting in K426R (transformylase region) and a frameshift in the other allele. This individual presented with profound developmental delay. It is likely that most mutations in ATIC are embryonic lethals, consistent with its critical role in DNPS and metabolism. Since ZMP is the substrate for ATIC, ATIC deficiency is likely to activate AMPK and have major consequences for cellular metabolism. DNPS nulls were recently generated in HeLa cells via CRISPR-Cas9 [16]. The crATIC cell line [16], which has no ATIC activity and accumulates ZMP, will allow detailed study of the consequences of mutations in ATIC. For example, given that adenine supplementation effectively shuts down DNPS [17,18], analysis of crATIC in adenine depleted growth media will allow detailed assessment of the effect of ZMP accumulation on transcription, translation, and metabolism. Transfection of crATIC with ATIC clones containing various mutations will allow cellular analysis of specific amino acid substitutions on ATIC structure and function. Clinical manifestations of DNPS deficiency have not been resolved through targeted therapeutics. This is likely due to an inadequate understanding of the processes affected by DNPS. The current work represents a continued effort to elucidate and categorize processes influenced or regulated by DNPS. Here, we present an analysis of the crATIC and HeLa cells in the presence or absence of adenine to identify the processes affected by DNPS and ATIC deficiency.

**Figure 1.**
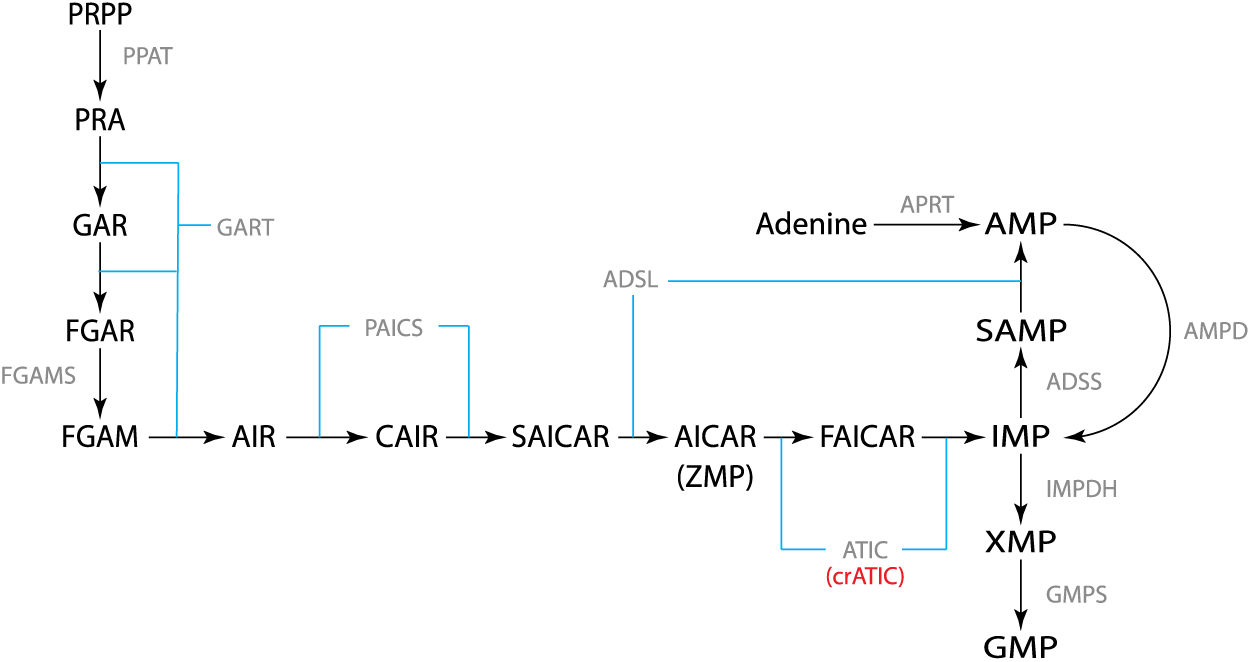
De novo purine synthesis pathway. DNPS is accomplished via the conversion of PRPP to IMP which is subsequently converted to AMP or GMP. The HeLa ATIC KO, crATIC, is indicated.

## 2. Materials and Methods

### 2.1. Cell Culture

crATIC cells were described previously [16]. HeLa cells were purchased from ATCC (Manassas VA USA). Cells were grown on 60 mm TPP (Techno Plastic Products, AG, Switzerland) plates using DMEM with 10% fetal calf serum (FCS), 30 μM adenine, and Normocin (InvivoGen). For purine depletion experiments (starvation conditions) cells were incubated in similar medium but using fetal calf macroserum (FCM: serum dialyzed against a 3.5 kDa barrier) with or without 100 μM adenine. Complete media was regularly refreshed, and ten to twelve hours before experiments, was replaced with adenine-depleted or 100 μM adenine-supplemented media. Cells were subjected to starvation conditions for ten hours. For histological staining, 10,000 cells were plated overnight in complete growth media (DMEM 10%FCS with 30 μM adenine and Normocin), media was replaced with starvation conditions (DMEM 10% FCM, Normocin, with or without 100 μM adenine) and after six days were fixed using 10% ethanol / 3.5% acetic acid solution and stained using 0.1% crystal violet. Entire culture areas were imaged.

### 2.2. HPLC analysis of ZMP metabolite accumulation

At two-hour intervals for ten hours, cells were washed once with cold 1x PBS and extracted with 500 μl ice cold 80% ethanol. Plates were scraped and the extract was centrifuged twice at 14,000 x g. Supernatant was collected and dried using a Speedvac then resuspended in 300 μl freshly prepared mobile phase. Samples were cleared twice by centrifugation at 14,000 x g and stored at -20 ºC until analysis. HPLC-EC separation of ZMP was as previously described [19]. Analytes were detected using a CoulArray HPLC system (model 5600A, ESA) with EC channel potentials set from 0 to 900 mV in 100 mV increments, then 1200 mV, and 0 mV. The autosampler was kept at 10 ºC over the course of the runs. Sum of primary peak area was used to assess analyte accumulation.

### 2.3. RNA-seq

Cells were cultured as described above. RNA-seq was performed as previously described [19]. Four replicates (a single culture was split into four plates) of crATIC and HeLa were cultured in adenine-supplemented or adenine-depleted (starvation) media for 10 hours. Technical replicates were used to ensure the accuracy in our measurements between the cell lineages and purine supplementation. Total RNA was extracted using Tri-reagent (Sigma) according to the manufacturer’s protocol, followed by spin columns (Machery Nagel) and elution with 50 μl DEPC treated water (Sigma). RNA purity and concentration were assessed using a NanoDrop One (Thermo Scientific). RNA was stored at -80 ºC. RNA quality assessment and RNA-seq was performed by The Genomics and Microarray Core Facility at the University of Colorado, Denver. mRNA libraries were constructed using the Nugen Universal Plus mRNA-Seq + UDI kit (cat # 9144–96), and 50 bp single read sequencing was performed employing the Illumina HiSEQ4000. The sequencing quality Q score was >38 for all reads. Conversion of .bcl to FASTQ files was done using CASAVA 2.0.

### 2.4. Data Processing

Computation was performed as previously described [19]. For each comparison group, the gene_exp.diff file (Cufflinks output) was filtered for significant entries where FPKM values ≥ 1 and log2 fold change values were log2 ≥ 1 or log2 ≤ -1. In each comparison group the 100 differentially expressed genes (DEGs) with the highest absolute log2 values (both 100 most positive and 100 most negative) were combined for ClueGO analysis. Total gene lists from our cutoffs were used for BiNGO analysis. Comparisons of crATIC to HeLa in adenine-depleted conditions are labeled as MM (minus to minus) or in adenine-supplemented conditions labeled as PP (plus to plus).

### 2.5. Gene Ontology analysis

Cytoscape (version 3.7.0) apps ClueGO (version 2.5.2) and BiNGO (version 3.0.3) were used to provide representative biological information from the lists of differentially expressed genes (DEGs). ClueGO using the top 100 most positive and 100 most negative DEGs was run using pairwise comparisons: HeLa plus adenine vs crATIC plus adenine (PP) and HeLa minus adenine vs crATIC minus adenine (MM). BiNGO is capable of utilizing our complete DEG lists but cannot run pairwise comparisons. The BiNGO analysis was limited to running one list of DEGs as a supplement to ClueGO results. The ClueGO analysis included biological process, cellular component, and molecular function gene ontologies as well as Reactome pathway and reactions, while BiNGO was only run with biological process, cellular component, and molecular function gene ontologies.

### 2.6. qPCR validation of DEGs

qPCR was performed to validate the RNA-seq analysis. Total RNA was isolated as described above with cDNA prepared using the iScript cDNA synthesis kit (Bio-Rad #1708890) and qPCR conditions previously described [19]. Candidate gene primers were purchased from IDT PrimeTime service: TGFβI (Hs.PT.58.40018323), ALPP (Hs.PT.56a.38602874.g), Twist1 (Hs.PT.58.18940950), IQGAP2 (Hs.PT.58.28018594), GATA3 (Hs.PT.58.19431110), β-Actin (Hs.PT.39a.22214847), OASL (Hs.PT.58.50426392), TUSC3 (Hs.PT.58.3740957), and DPYSL3 (Hs.PT.58.39796068). C_t_ values were normalized to β-Actin.

## 3. Results

### 3.1. crATIC requires adenine for proliferative growth

HeLa and crATIC cells were cultured in complete media then subsequently cultured in DMEM supplemented with dialyzed fetal calf macroserum with or without adenine. HeLa cells showed proliferative growth in both media conditions while crATIC cells only showed proliferative growth in adenine-supplemented conditions (Figure 2).

**Figure 2.**
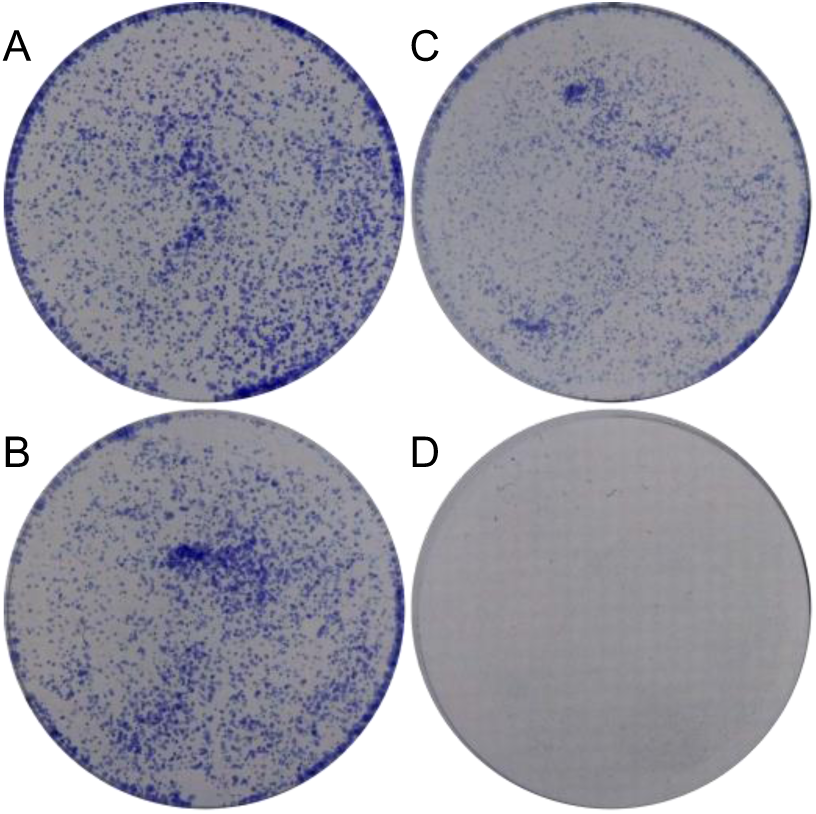
crATIC requires adenine for proliferative growth. HeLa (A,B) and crATIC (C,D) cells were plated and cultured in adenine-supplemented (A,C) or adenine-depleted (B,D) media. Plates were stained with crystal violet.

### 3.2. crATIC cells accumulate ZMP in adenine-depleted but not adenine-supplemented growth conditions

crATIC cells were cultured in adenine-depleted and adenine-supplemented media. Metabolites were ethanol extracted from cells in culture every two hours for ten hours. In samples from adenine-depleted (starved) cells, a ZMP peak was observed at the first time point in HPLC-EC traces (24.0 mins) and this peak increased linearly until the last time point measured (Figure 3 and Supplementary Figure 1). A ZMP peak was not observed in samples from adenine-supplemented cells, which is consistent with previous results [19]. Ten hours of starvation was chosen for experimentation moving forward. This is due to a few reasons: 1) prior experimentation from our lab used a ten hour time point for the characterization of the crADSL cell line [19] 2) the DNPS-KO cell line begins to show an unhealthy phenotype within 2-3 days of starvation, and therefore 3) this time point likely balances accumulation and transcriptional changes in a healthy cell versus the time of induction to cell death signaling or cell cycle stalling process. DNPS is upregulated at the G1/S phase interface with ten hours representing approximately 40% of the cell cycle in HeLa cells.

**Figure 3.**
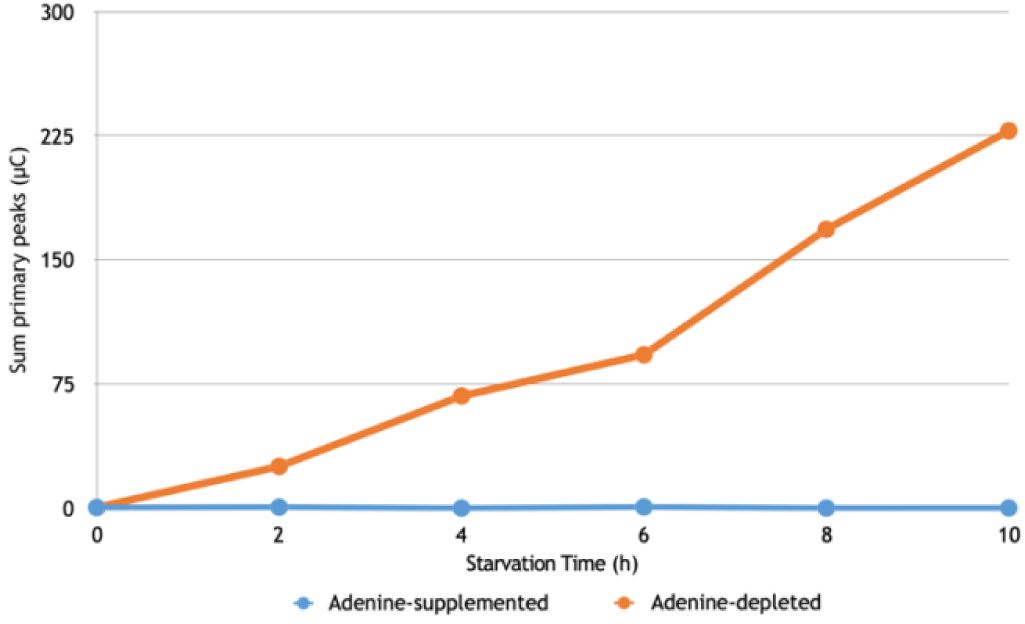
ZMP accumulates in crATIC. crATIC was grown in media with or without supplemental adenine. Accumulation of ZMP was measured by HPLC-EC and was observed only in cells grown in adenine-depleted media.

### 3.3. crATIC shows differentially expressed genes compared to HeLa in adenine-depleted and adenine-supplemented conditions

The crATIC and HeLa cell transcriptomes were compared after culture in adenine-supplemented (PP) and adenine-depleted (MM) conditions. In both comparison groups, DEGs were limited by cutoffs (see Materials and Methods). In the PP comparison, we obtained 1311 DEGs while in the MM comparison, we obtained 1662 DEGs. This suggests that adenine supplementation corrects for some of the differences in gene transcription in the two cell types. The DEGs with the top 100 (positive) and bottom 100 (negative) log2 values were selected for further analysis. In the PP comparison, these represent log(2)fold ranges of -8.60573 to -2.31272 and 2.26702 to 8.20331. In the MM comparison, the DEGs with the top 100 positive and bottom 100 negative log2 values represent log(2)fold ranges of 8.39229 to 2.31793 and -8.87985 to -2.41031. These 200 gene lists were used for ClueGO analysis (Figure 4).

**Figure 4.**
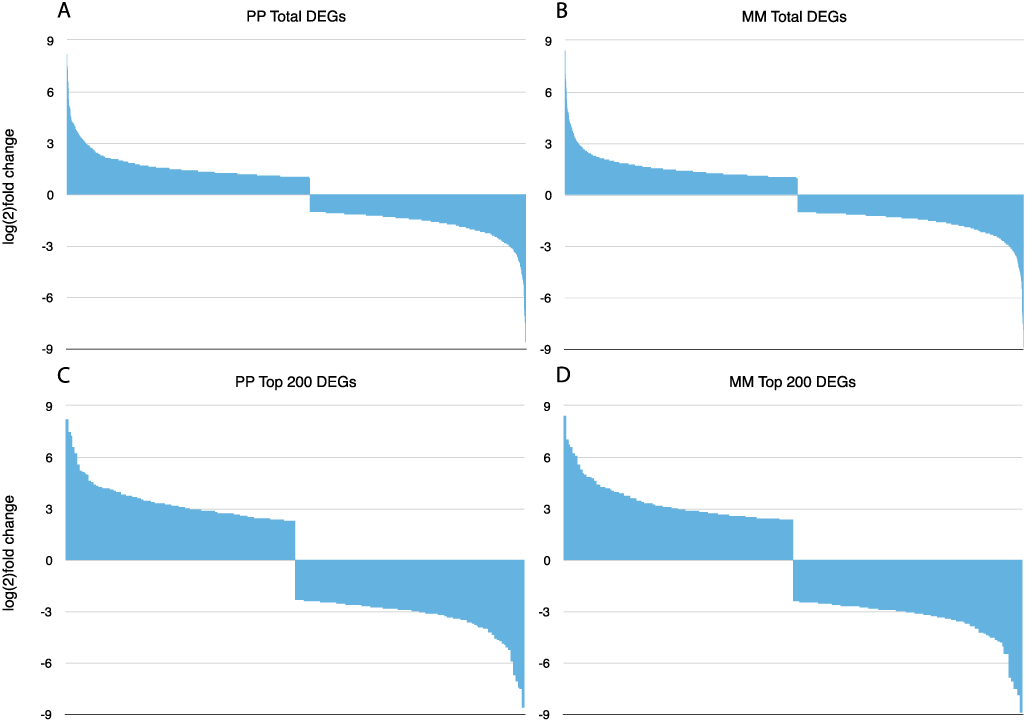
log(2) fold change of DEGs in cell lines under experimental conditions. A and B: DEGs in PP (A) and MM (B) conditions. C and D: 100 most positively and 100 most negatively changed DEGs in PP (C) and MM (D) conditions.

Comparing the MM and PP groups, 228 genes were unique to the PP group while 579 genes were unique to the MM group. 1083 genes were common to both groups. When comparing DEGs with the 100 most positive and 100 most negative log2 values (200 total), 44 genes were found unique to each group, and 156 genes were shared between the two groups (Table 1). Principle component analysis showed tight clustering by cell type and media supplementation (Figure 5).

**Table 1.** Number of unique and shared DEGs

|  | Total DEGs | Top 200 DEGs |
| --- | --- | --- |
| Shared | 1083 | 156 |
| Unique PP | 228 | 44 |
| Unique MM | 579 | 44 |

**Figure 5.**
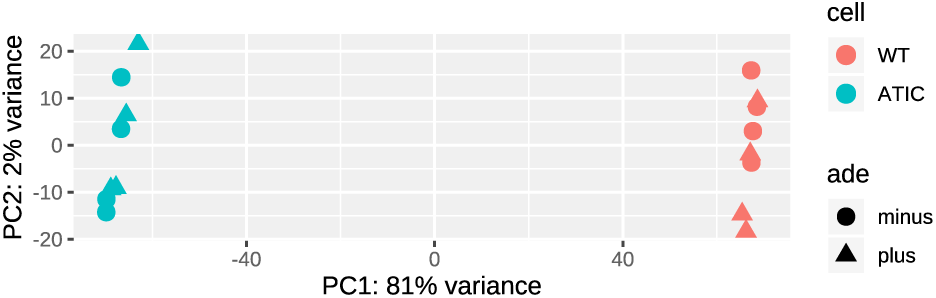
Principle component analysis of crATIC and HeLa experimental groups. PCA shows robust differences by cell type.

### 3.4. DEGs showed enrichment in ClueGO Gene Ontology and Reactome database analysis

RNA-seq experiments can provide important data bearing on global cellular changes in response to stressors and changes in environment.

The lists of 200 DEGs described above were used as input for ClueGO, a Cytoscape application that extracts representative functional biological information for large lists of genes or proteins [20,21]. We queried Gene Ontology (GO, biological process, cellular component, and molecular function [22]) and the Reactome Knowledgebase (pathways and reactions [23]) to identify significant terms from comparison of the crADSL and HeLa transcriptomes. Although we identified many GO and Reactome terms (see Supplementary Information), we focused on an interesting subset of specific terms implicated in Alzheimer’s disease and cellular aging.

For Biological Process GO ontologies, we obtained 55 groups: 34 associated with shared terms, 33 associated with the PP comparison, and 23 associated with the MM comparison (prominent GO groups are shown in Figure 6). Notable shared terms included inflammation / immune response terms such as IL-1β secretion, t-cell activation involved in immune response, eicosanoid metabolic process, prostanoid metabolic process, prostaglandin metabolic process, negative regulation of TNF product, response to amyloid beta, glutamine family biosynthetic process, and long chain fatty acid biosynthetic process and transport. Regulation of pathway-restricted SMAD protein phosphorylation and transforming growth factor beta terms were also in the list. For the PP comparison group there was enrichment of fatty acid terms, including arachidonic acid metabolic process, and the cyclooxygenase pathway. Other terms included neuron cellular homeostasis and neuron migration, midbrain development, and neurotransmitter metabolic process. Regulation of cellular response to TGFβ stimulus, response to hyperoxia, negative regulation of embryonic development, and negative regulation of G1/S transition of mitotic cell cycle were also enriched terms. For the MM comparison, neuroinflammatory response, regulation of epithelial to mesenchymal transition, negative regulation of erk1/2 cascade, cellular response to fatty acid, and cellular response to prostaglandin stimulus were obtained in the analysis (Supplementary Figures 2 and 3).

**Figure 6.**
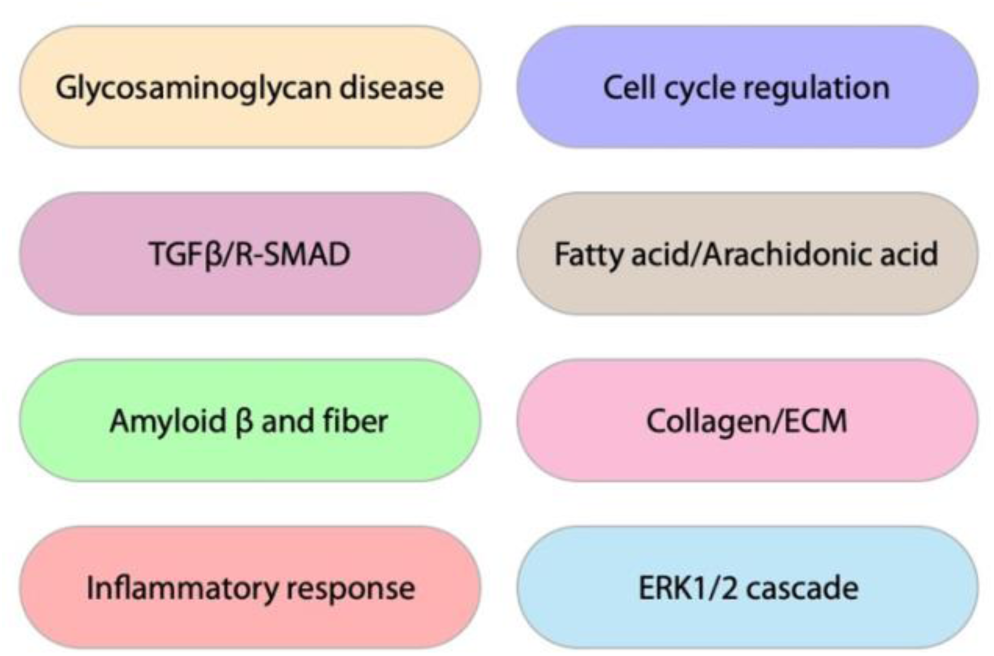
Notable GO groups from ClueGO analysis

For Cellular Component ontologies, we obtained 10 groups. Five groups were from shared comparisons, 2 from PP and 5 from MM. Shared terms included golgi lumen, melanosome, and complex of collagen trimers. Terms enriched in the PP comparison included dystrophin-associated glycoprotein complex and brush border. Terms associated with the MM comparison included extracellular matrix component, cell division site, and actomyosin (Supplementary Figures 4 and 5).

In the Molecular Function ontologies, 14 groups of terms were identified with 5 groups from shared, 6 groups from the PP comparison, and 3 from the MM comparison. Terms found within the shared groupings were collagen binding, proteoglycan binding, neuropeptide receptor binding, and amino acid binding. In the PP comparison, terms of note include steroid hormone receptor activity, adenylyl transferase activity, cell-cell adhesion mediator activity, and lysophospholipase activity. In the MM comparison, terms such as Aβ binding and l-amino acid transmembrane transporter activity were found (Supplementary Figures 6 and 7).

**Figure 7:**
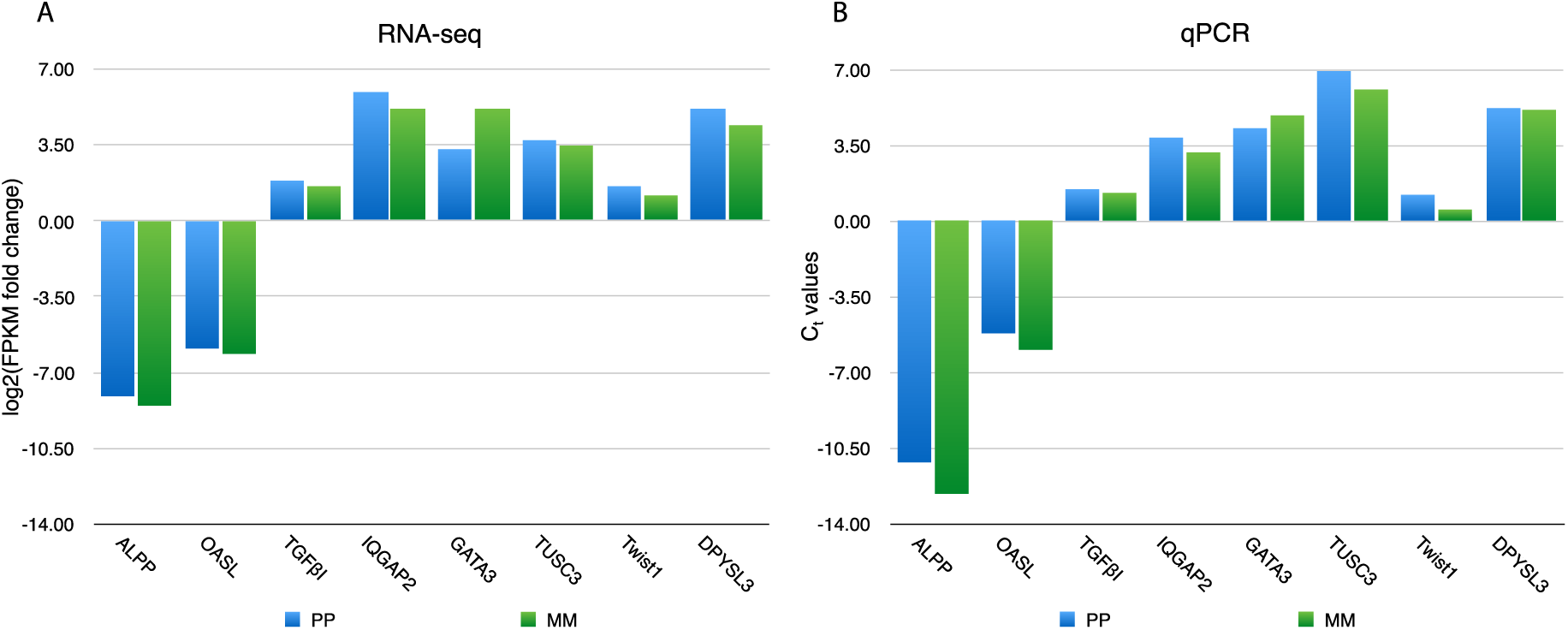
Candidate gene verification by qPCR. Gene transcription in PP and MM conditions measured by A) log(2)fold change of FPKM values from RNA-seq and B) Ct values from qPCR. Values were normalized to β-Actin.

In the results from querying the Reactome Knowledgebase, we obtained 10 groups in Pathways, 5 shared, 6 from the PP comparison, and 2 from the MM comparison. Within the shared groupings, we obtained terms such as signaling by retinoic acid, amyloid fiber formation, signaling by NOTCH1, arachidonic acid metabolism, and collagen formation. In the PP comparison we obtained nuclear receptor transcription pathway, TP53 regulates transcription of cell cycle genes, transcriptional regulation of the AP-2 (TFAP2) family of transcription factors, IL-4 and IL-13 signaling, and synthesis of prostaglandins (PG) and thromboxanes (TX). In the MM comparison we obtained IL-10 signaling, ECM proteoglycans, and diseases associated with glycosaminoglycan metabolism (Supplementary Figures 8 and 9).

In Reactome Reactions, we obtained 7 groups of terms with 4 groups shared, 3 groups from the PP comparison, and no groups from the MM comparison. Shared groups included expression of IFNγ-stimulated genes, exocytosis of specific granule membrane proteins, and terms associated with collagen. In the PP comparison we obtained formation of NR-MED1 coactivator complex, keratin filaments bind cell-cell adhesion complexes, and binding of AP1 transcriptional activator complexes to CCND1 promoter (Supplementary Figures 10 and 11).

### 3.5. Corroboration of ClueGO results via BiNGO analysis

To validate our ClueGO results, we performed enrichment analysis using the BiNGO application in Cytoscape [24] and the complete significant DEG lists. While we obtained some new terms, on the whole this analysis produced term enrichment closely similar to that from our ClueGO analysis (Supplementary Figures 12-17). Full tables of DEGs, ClueGO, and BiNGO results are available in Supplementary Tables 1-3.

### 3.6. Validation of RNA-seq by qPCR of candidate genes

To assess the validity of our RNA-seq results, we employed qPCR to confirm transcript levels for select genes. The results for these genes showed similar transcript levels and trends based on C_t_ value comparison with the log(2)fold values from our RNA-seq results (Figure 7).

## 4. Discussion

In the present study, we evaluated dietary requirements for crATIC and metabolite accumulation during purine starvation.

In addition to its role in rescuing the DNPS KO phenotype, adenine supplementation shuts down DNPS [17,18] and prevents ZMP accumulation in crATIC. Adenine phosphoribosyltransferase (APRT) mediates conversion of adenine and phosphoribosyl pyrophosphate (PRPP) to AMP and pyrophosphate (PPi) and is present in all mammalian tissues. APRT is uniquely responsible for metabolic adenine salvage from dietary sources [25]. Adenine supplementation also provides a source for synthesis of GMP. AMP can be converted to IMP by AMP deaminase, and then converted to GMP by IMP dehydrogenase [IMPDH, converts IMP into xanthine monophosphate (XMP)] and GMP synthase (converts XMP into GMP) [26].

We employed RNA-seq to compare the crATIC and HeLa transcriptomes in adenine supplemented and adenine-depleted conditions, and we performed qPCR to verify these results. We established a 10-hour starvation (cell culture in adenine-depleted media). At the end of this time course, ZMP levels were still increasing in crATIC cells, and we have not yet established a time point at which ZMP levels plateau. It is possible ZMP accumulation during this period may not have been adequate for full AMPK activation. As discussed above, ZMP is an AMP mimetic and likely has multiple enzymatic targets. The crATIC HeLa system can be employed to identify these targets.

Our results demonstrate that crATIC requires purine (adenine) supplementation for proliferative growth, and that ZMP accumulates linearly over a time course of ten hours during purine starvation. After analysis of the RNA-seq data, we obtained many DEGs both by cell type and adenine supplementation. HeLa cells were chosen due to current availability [16] and their potential for broad scale information about cellular processes affected by DNPS deficiency and translation to disease states.

Although DEGs mapped to numerous gene ontologies, we focused our discussion on terms of specific interest. AMPK sensitivity is decreased in aged models [27], however in Alzheimer’s disease (AD), an aging related disorder, pAMPK is abnormally elevated in tangle and pre-tangle bearing neurons [28]. Data also suggest that insulin resistance seen in AD reduces the astrocytic energy supply via inhibition of AMPK, contributing significantly to the neurodegeneration observed in AD [29]. AMPK has a major role in Tau protein phosphorylation, as well as mTOR and autophagy pathways, which are dysfunctional in AD [30]. Energy homeostasis in Alzheimer’s disease is also dysregulated possibly via an AMPK related mechanism [31], illustrating the need to understand energetic processes in dementia. Our results suggest a role for ZMP or ATIC in AD and possibly other amyloid or tau-based dementias. AD is a progressive, age related, dementia characterized by amyloid-β plaques, tau fibril formation, and extensive neuronal degradation in the brain. One feature of AD is chronic neural inflammation [32]. Typically, inflammation is caused by a cellular response to stimuli and is normally terminated by a process called resolution. Arachidonic acid (AA) may play an important role in the inflammatory response in AD [33], especially in initiation of resolution [34-36]. Chronic inflammation in AD is marked by a dysregulation of resolution in which the inflammatory response persists [34].

Normally, AA is a constituent of phospholipids. The circulating form is rare, as AA is rapidly scavenged and bound to albumin. Free AA can be cleaved from phospholipids by phospholipase (PLA) and is typically produced in response to injury or other stimuli. Free AA is metabolized rapidly through enzymatic pathways based on PLA protein-protein interactions with cyclooxygenase (COX), lipoxygenase (LOX), cytochrome P450 (CYP), fatty acid amide hydrolase (FAAH), or non-enzymatic lipid peroxidation and oxidative stress. FAAH is a concentration-dependent reversible reaction, producing anandamide (a neurotransmitter) from AA and ethanolamine, and AA release can be stimulated by anandamide [37,38]. AA metabolic products have roles in processes such as platelet aggregation, vasoconstriction and dilation, toxic shock based organ dysfunction, inflammatory response, female fertility, fever mediation, Alzheimer’s and Parkinson’s disease neurodegeneration, mediation of cAMP, suppression of excitatory neuronal signals via endocannabinoids, and smooth muscle response [39].

Cytosolic phospholipase A2 (cPLA2) is expressed in higher levels in brain tissue from AD patients [40] and eicosanoids, a product of AA COX and LOX based metabolism, act as mediators of neuroinflammation. In AD, COX2 is overexpressed, and this overexpression has been correlated with AD progression [41]. In addition, the COX1 splice variant, COX3, has been found in AD brain [42]. The prostaglandin metabolite, PGD2, is overproduced in glial cells surrounding amyloid plaques [43]. However, NSAIDs which act as COX2 inhibitors have so far been ineffective at mitigating the AD-associated inflammation [44]. Interestingly, LXA4, a metabolite produced via the LOX pathways from AA, has been implicated in reduction of reactive oxygen species [45], inhibition of interleukin expression [46] and reduction of Aβ levels [47]. LXA4 is also implicated in resolution of inflammation [48]. In cerebrospinal fluid and brain tissue from AD patients, LXA4 levels are lower than from control groups, while 15-LOX-2, the enzyme that catalyzes production of LXA4, was elevated in glial cells from AD patients [34]. These results suggest that AA metabolism is differentially regulated in AD. Our results suggest that ZMP, ATIC, and DNPS may have important roles in AA metabolism and may be attractive targets for further investigation and development of new therapies.

Microglial cells respond to diverse cues from injured neurons by becoming activated and inducing phagocytosis to initiate clearance of apoptotic cells or extruded proteins (e.g., Aβ). Activation is accompanied by release of cytokines such as IL-1β and TNFα [49] and regulatory cytokines [50]. There are inherent differences between aged, chronically activated microglia compared to young microglia [51]. In young microglia, cells are sensitive to TGFβ signaling which leads to a reduction in cytokine release [52], neuroprotection [53], and promotion of phagocytosis [54]. However, in chronically activated microglia, there is decreased phagocytosis [55], and a reduced response to TGFβ signaling, resulting in increased neurotoxicity and reduced uptake of Aβ [56,57]. This diminished response is largely attributed to reduced TGFβ-Smad coupling, the canonical pathway of TGFβ signaling, that occurs in AD [58] and aging [59]. In non-canonical pathways, TGFβ activates MAPK, PI3K, and JNK: each are linked under certain situations to a pro-inflammatory response [60]. In canonical TGFβ signaling, activation of the TGFβ-Smad pathway induces glial cells to transcribe MAPK phosphatase (MKP), a serine-threonine phosphatase responsible for negative regulation of inflammation that results in decreased TNFα production and reduced Aβ–mediated activation of MAPK and NFκB signaling [61]. However, in cases of chronic microglial inflammation observed in aging and AD, the TGFβ-Smad pathway is inhibited, resulting in reduced amelioration of inflammation. Our identification of ATIC and DNPS involvement in TGFβ signaling coupled with FA based inflammatory regulation provides a novel avenue for continued AD research.

During embryogenesis and fetal development, purines have an important role as signaling molecules [62], particularly in neuronal development [63-65]. DNPS has been found upregulated at the G1/S cell cycle transition [66,67]. So far, mutations that cause reduction in PAICS, ADSL, or ATIC activity in humans all lead to significant developmental delay. These findings indicate the importance of the pathway for producing purines for DNA and RNA synthesis in embryogenesis and development. Consistent with these roles, in our comparison of the crATIC and HeLa transcriptomes, we identified many DEGs that mapped to ontology terms concerned with development and neuronal function as well as ontology terms associated with rapid cellular proliferation, such as tumorigenesis and cancer. Multiple terms also mapped to cell cycle checkpoints and terms associated with the G1/S phase interface.

Previously, we characterized the crADSL cell line transcriptome [19]. Common ontology terms from the crADSL and crATIC transcriptome analyses are likely related to general DNPS function rather than the specific mutated DNPS enzyme or accumulated intermediates. Common terms include transforming growth factor beta, collagen/ECM, glycosaminoglycan metabolism, interferons, and embryonic development. Terms specific to the crATIC analysis include G1/S checkpoint transition, dystrophin associated glycoprotein complex, retinoic acid response, arachidonic acid metabolism, and prostaglandin and thromboxane. Of note, phospholipase activity was common to both crADSL and crATIC while lysophospholipase was found only in the crATIC analysis.

As discussed above, the crATIC cell line is advantageous in that ZMP accumulation is achieved via cellular metabolism, not conversion of exogenous AICAr in the media. The crATIC cells and crATIC cells transfected with identified patient allele and other mutant forms of ATIC should provide important cellular models of DNPS, AICA-ribosiduria, and ZMP accumulation. The results reported here are an important first step in establishing this model, and for further investigation of ZMP’s role as an AMP mimetic.

## Supporting information

Supplementary Table 1 Genes list

Supplementary Table 2 ClueGO

Supplementary Table 3 BiNGO

## Acknowledgements

The University of Colorado Cancer Center and the Genomics (Microarray) Shared Resource are supported in part by the Cancer Center Support Grant #P30-CA046934 from the National Cancer Institute.

MZ and VB were supported by Charles University [programmes PRIMUS/17/MED/6 and PROGRES Q26/LF1] and by the Ministry of Education, Youth and Sports of CR [LQ1604 National Sustainability Programme II].

## Author Contributions

R.M., D.P., and G.V. designed the experiments. R.M. performed ClueGO and BinGO analyses, qPCR validation of RNA-seq results, and was primary author of the manuscript. V.B. and M.Z. created and provided the CRISPR-Cas9 ATIC KO HeLa cell line, crATIC. N.D. performed HPLC-EC analysis to detect DNPS metabolites and intermediates. T.G.W. provided sample preparation expertise. G.V. performed salmon and DESeq2 processing of fastq sequence and provided informatics support. G.V. and D.P. directed and supervised the team members and manuscript preparation. All authors reviewed and edited the manuscript.

## Abbreviations

(APRT): adenine phosphoribosyltransferase
(AMP): adenosine monophosphate
(AMPK): AMP-activated protein kinase
(ATP): adenosine triphosphate
(ADSL): adenylosuccinate lyase
(AICAr): 5-aminoimidazole-4-carboxamide ribonucleoside
(ZMP): 5-aminoimidazole-4-carboxamide ribonucleotide
(ATIC): 5-aminoimidazole-4-carboxamide ribonucleotide formyltransferase / inosine monophosphate cyclohydrolase
(AA): arachidonic acid
(COX): cyclooxygenase
(CYP): cytochrome P450
(cPLA2): cytosolic phospholipase A2
(DNPS): *de novo* purine synthesis
(DEG): differentially expressed gene
(FDR): false discovery rate
(FAAH): fatty acid amide hydrolase
(FCM): fetal calf macroserum
(FCS): fetal calf serum
(FAICAR): 5-formamido-4-imidazolecarboxamide ribonucleotide
(FPKM): fragments per kilobase of exon per million reads mapped
(GO): gene ontology
(GMP): guanosine monophosphate
(IMP): inosine monophosphate
(INF): interferon
(LOX): lipoxygenase
(mTOR): mammalian Target of Rapamycin
(MM): minus adenine crADSL to minus adenine WT comparison
(PLA): phospholipase
(PAICS): phosphoribosylaminoimidazole carboxylase / phosphoribosylaminoimidazole succinocarboxamide synthetase
(PRPP): phosphoribosyl pyrophosphate
(PP): plus adenine crATIC to plus adenine WT comparison
(TSC1 and TSC2): Tuberous Sclerosis Complex 1 and 2
(XMP): xanthine monophosphate

