## Supplementary Figures 1-17 for "The CRISPR-Cas9 crATIC HeLa transcriptome: Characterization of a novel cellular model of ATIC deficiency and ZMP accumulation"

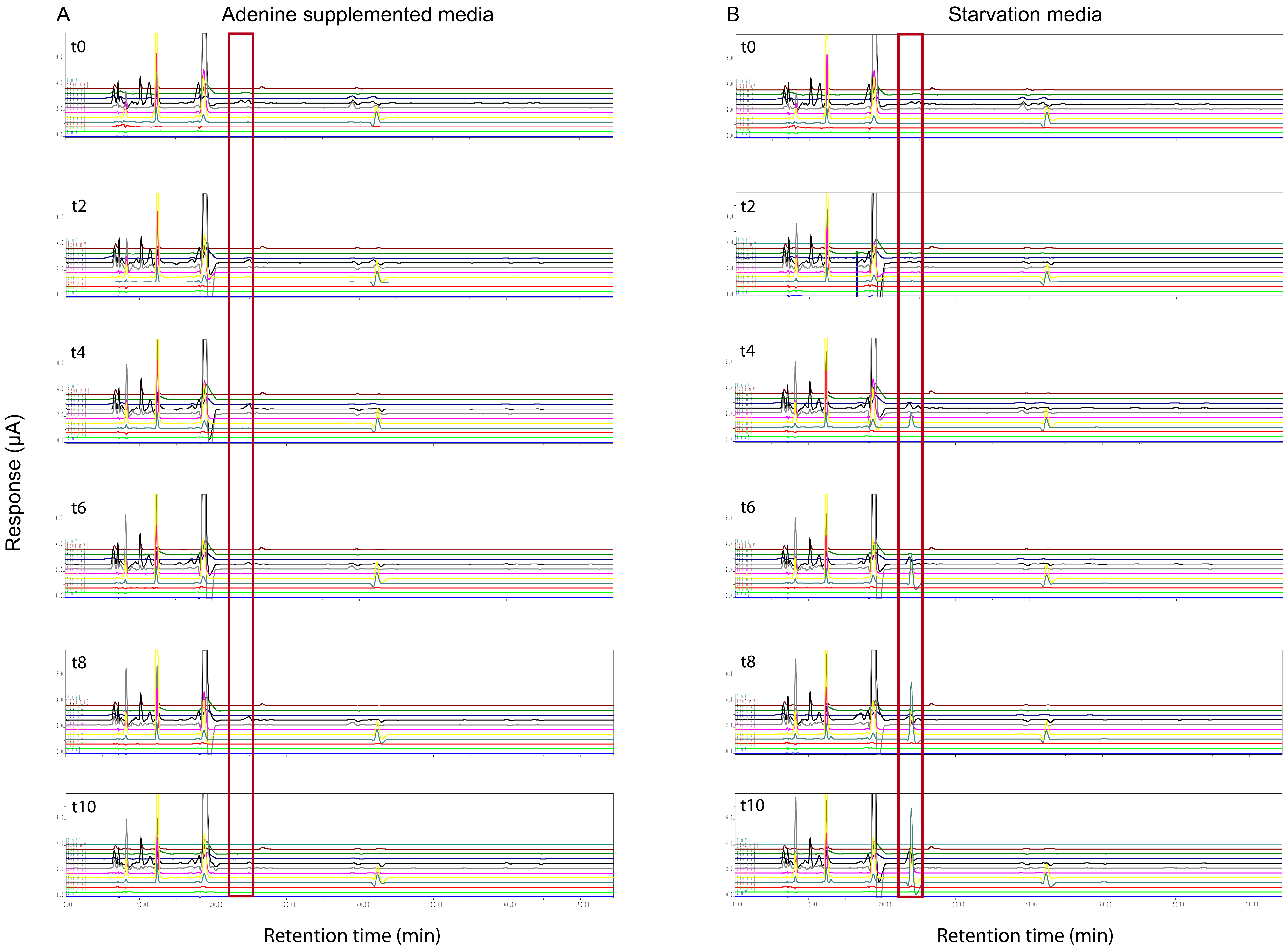


**Supplementary Figure 1: HPLC-EC traces of metabolites from crATIC cells.** Cells were cultured in FCM supplemented with adenine (A) and without adenine (B) for two-hour increments. The red rectangle indicates the elution time of ZMP with primary peak in channel 4 (300mV). C: Aligned and cropped t2 chromatogram: supplemented (plus), unsupplemented (minus) or unsupplemented and spiked (minus spike). The dashed line indicates the primary channel peak height (note starved sample and starved sample with ZMP spike).

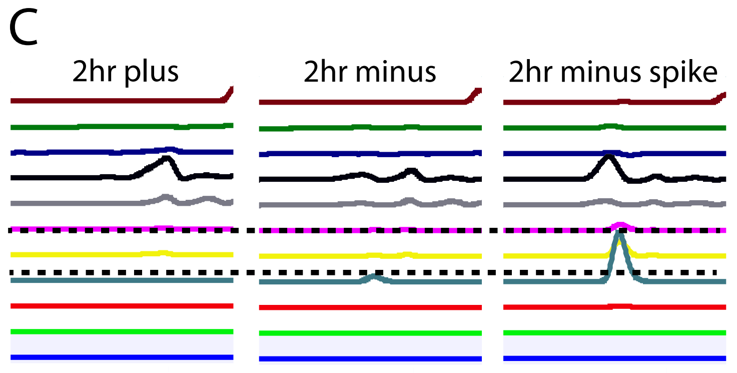

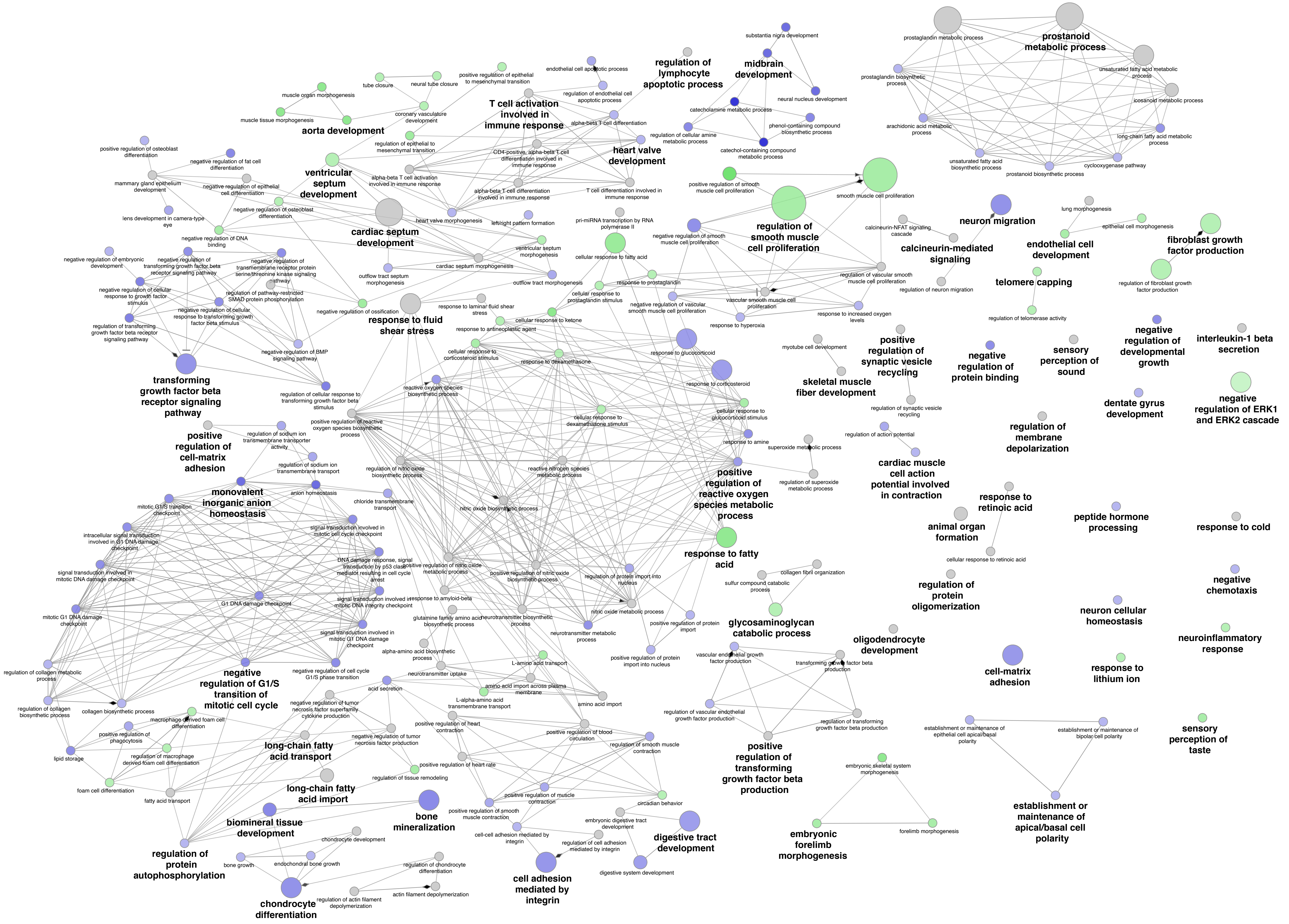


**Supplementary Figure 2: ClueGO Biological process network map.** Grey nodes indicate shared terms, blue nodes indicate terms enriched in PP comparison and green nodes indicate terms enriched in MM comparison.

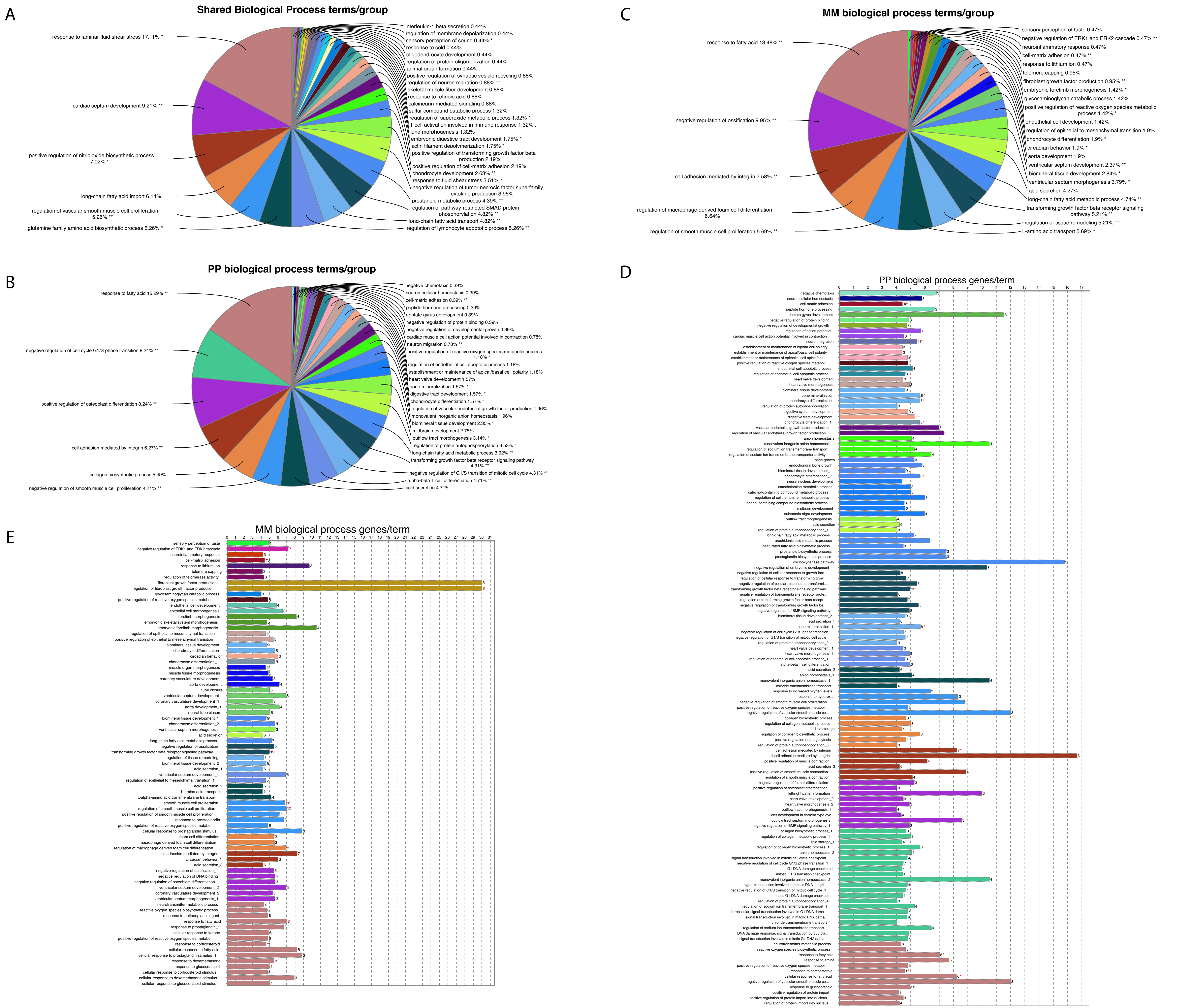


**Supplementary Figure 3: ClueGO Biological process terms and groups.** Pie charts show the proportion of the terms associated with the groups for Shared (A), PP (B), and MM (C). Histograms indicate number of genes found per term for PP (D) and MM (E) comparisons.

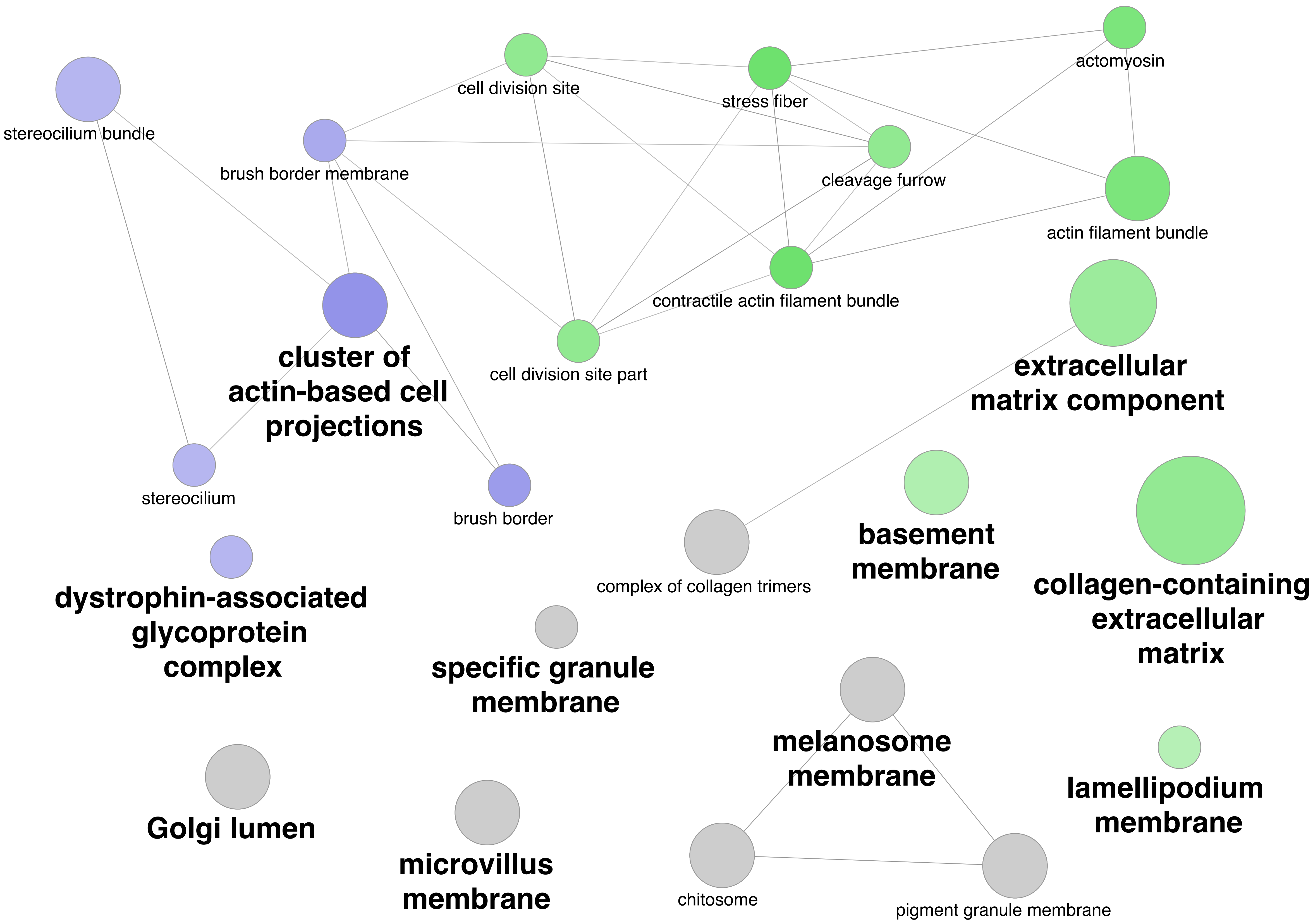


**Supplementary Figure 4: ClueGO Cellular component network map.** Grey nodes indicate shared terms, blue nodes indicated terms enriched in PP comparison, and green nodes indicate terms enriched in MM comparison.

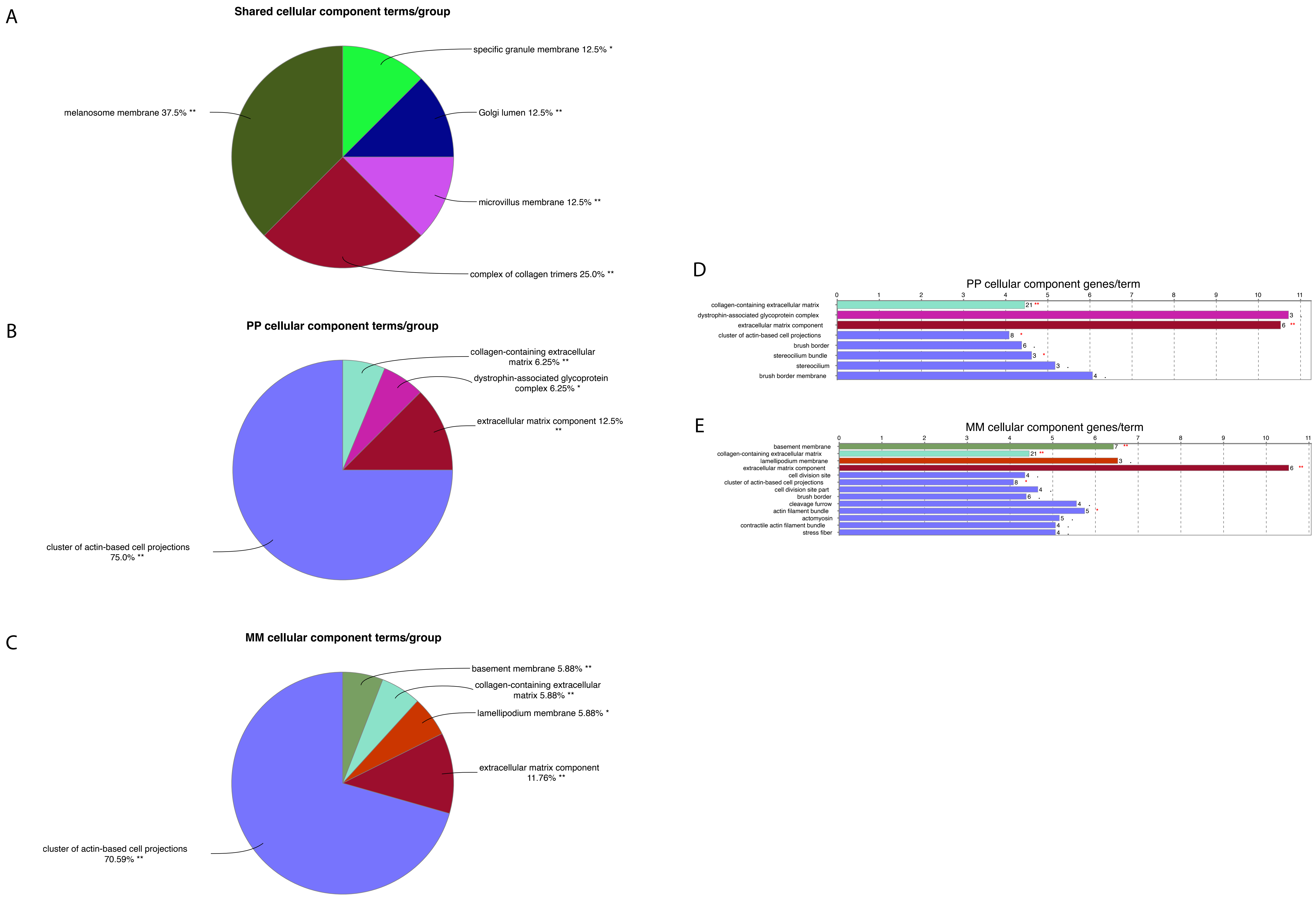


**Supplementary Figure 5: ClueGO Cellular component terms and groups.** Pie charts show the proportion of the terms associated with the groups for Shared (A), PP (B), and MM (C). Histograms indicate number of genes found per term for PP (D) and MM (E) comparisons.

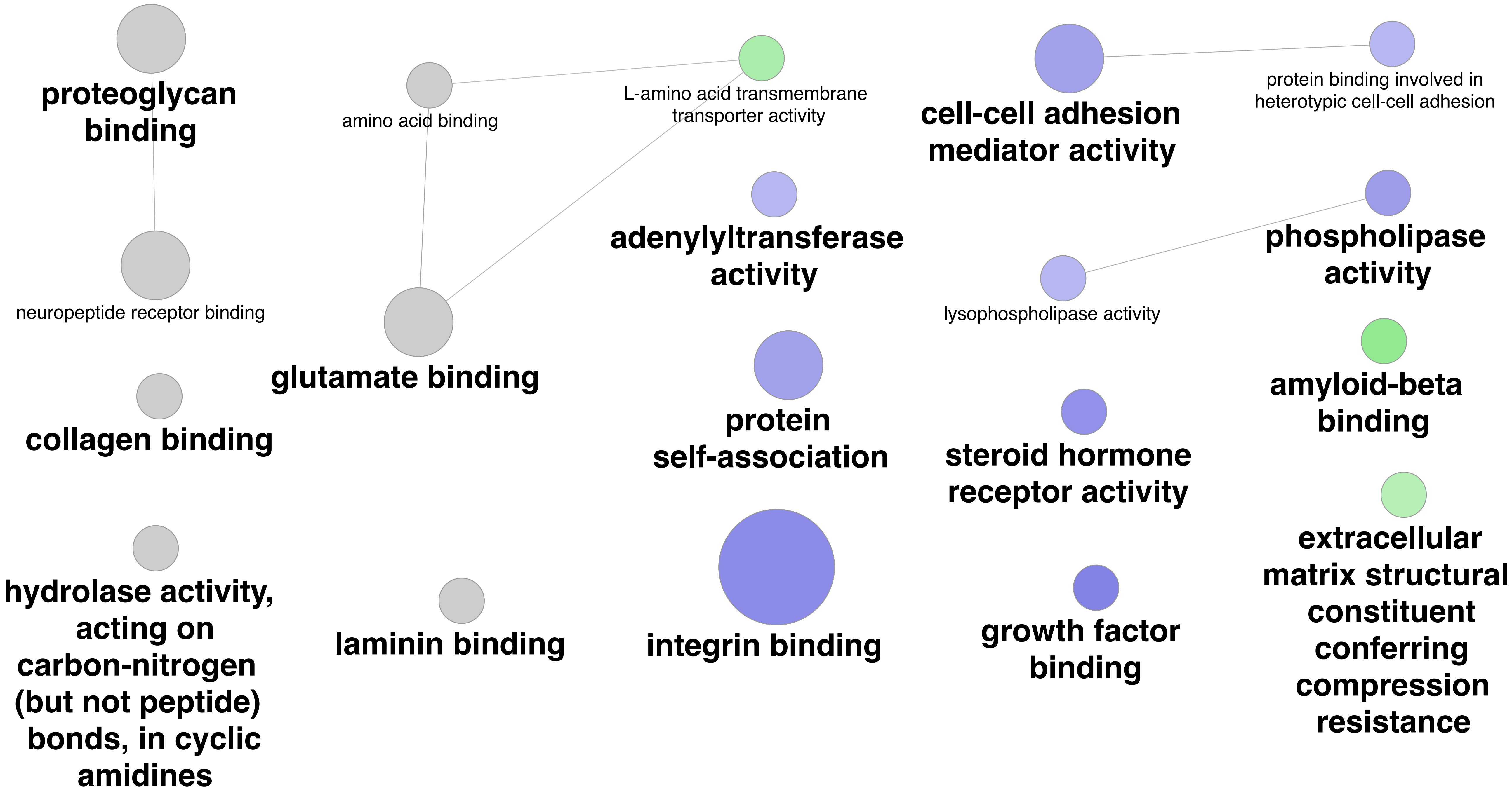


**Supplementary Figure 6: ClueGO Molecular function network map.** Grey nodes indicate shared terms, blue nodes indicated terms enriched in PP comparison, and green nodes indicate terms enriched in MM comparison.

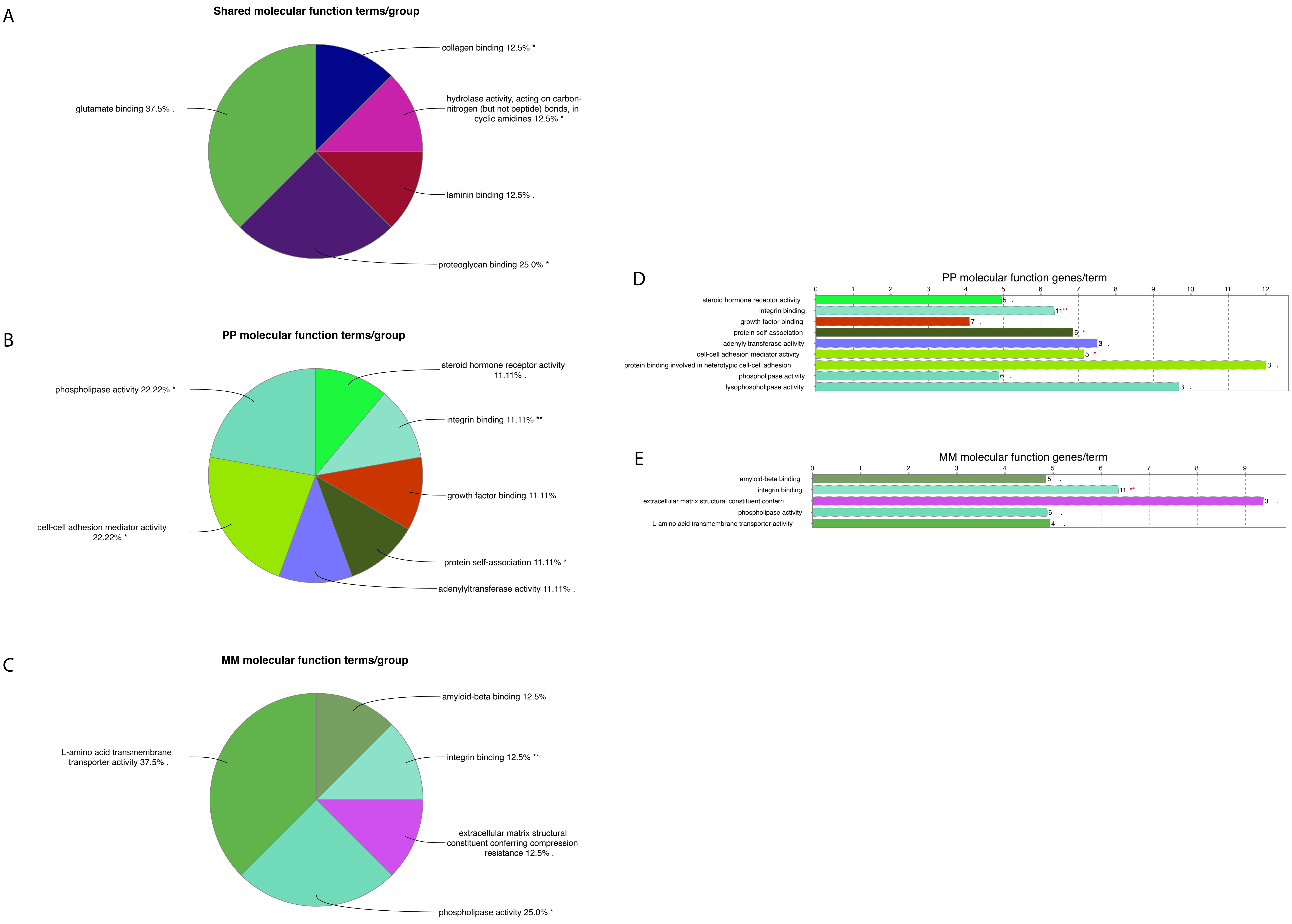


**Supplementary Figure 7: ClueGO Molecular function terms and groups.** Pie charts show the proportion of the terms associated with the groups for Shared (A), PP (B), and MM (C). Histograms indicate number of genes found per term for PP (D) and MM (E) comparisons.

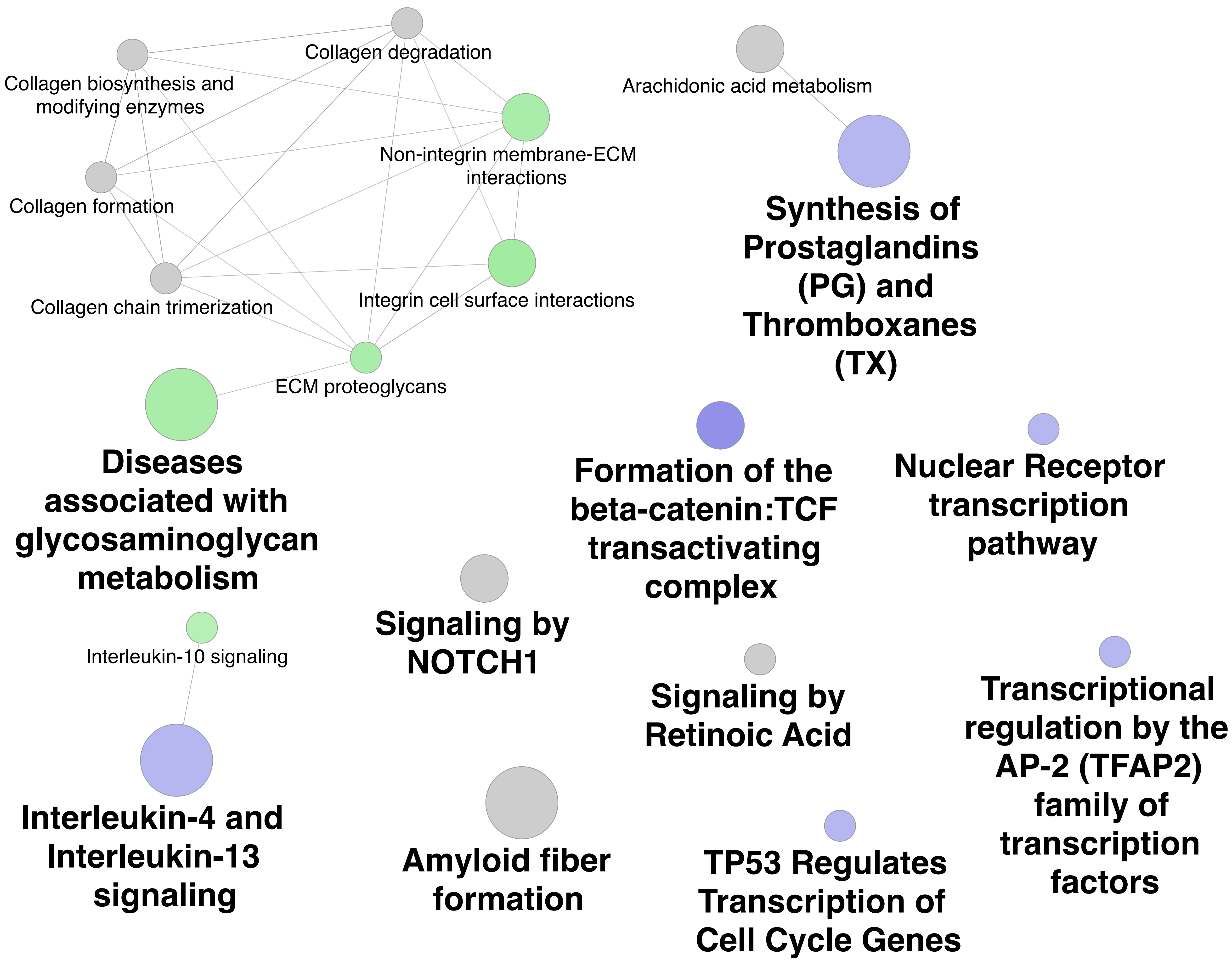


**Supplementary Figure 8: ClueGO Reactome pathways network map.** Grey nodes indicate shared terms, blue nodes indicated terms enriched in PP comparison, and green nodes indicate terms enriched in MM comparison.

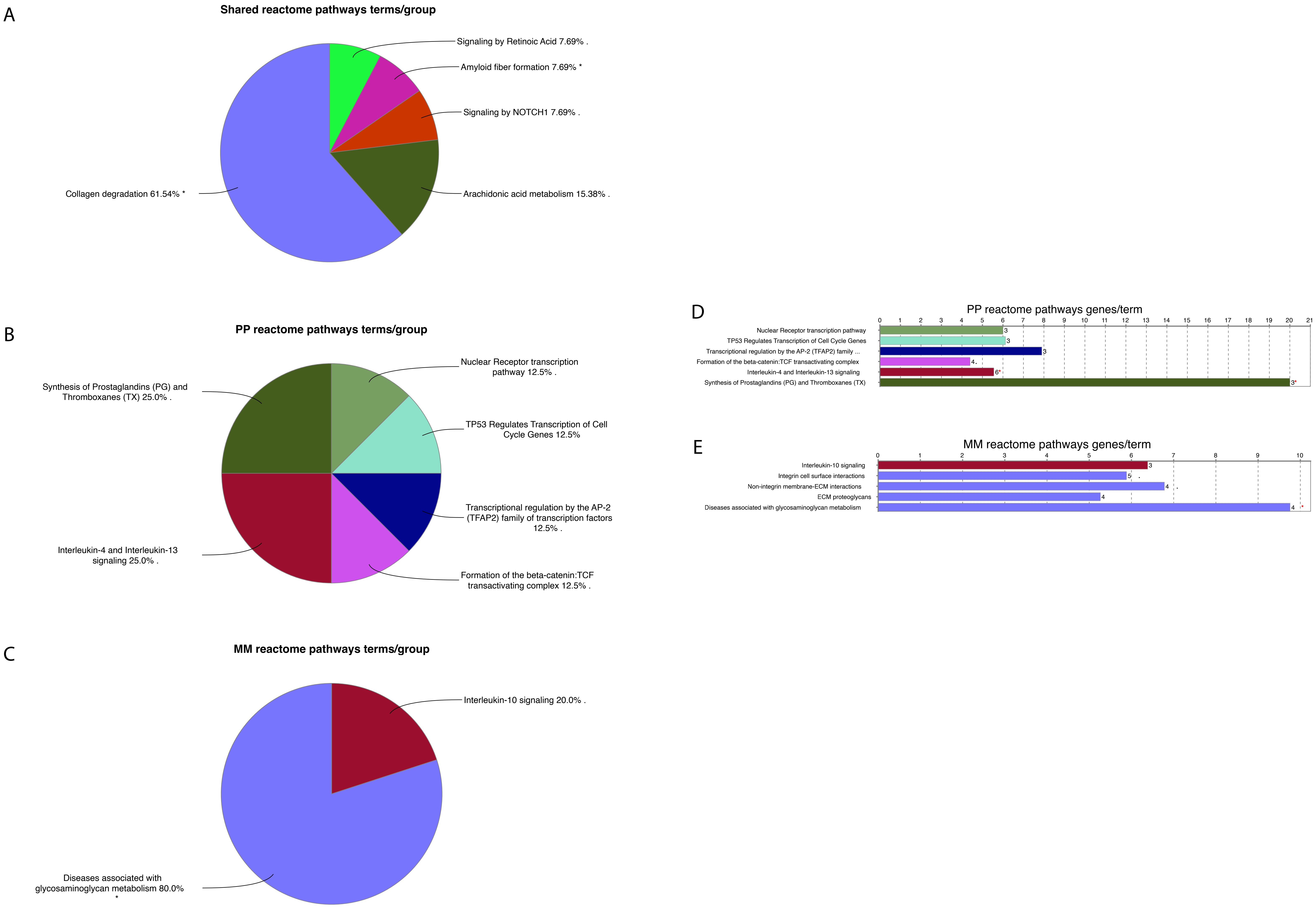


**Supplementary Figure 9: ClueGO Reactome pathways terms and groups.** Pie charts show the proportion of the terms associated with the groups for Shared (A), PP (B), and MM (C). Histograms indicate number of genes found per term for PP (D) and MM (E) comparisons.

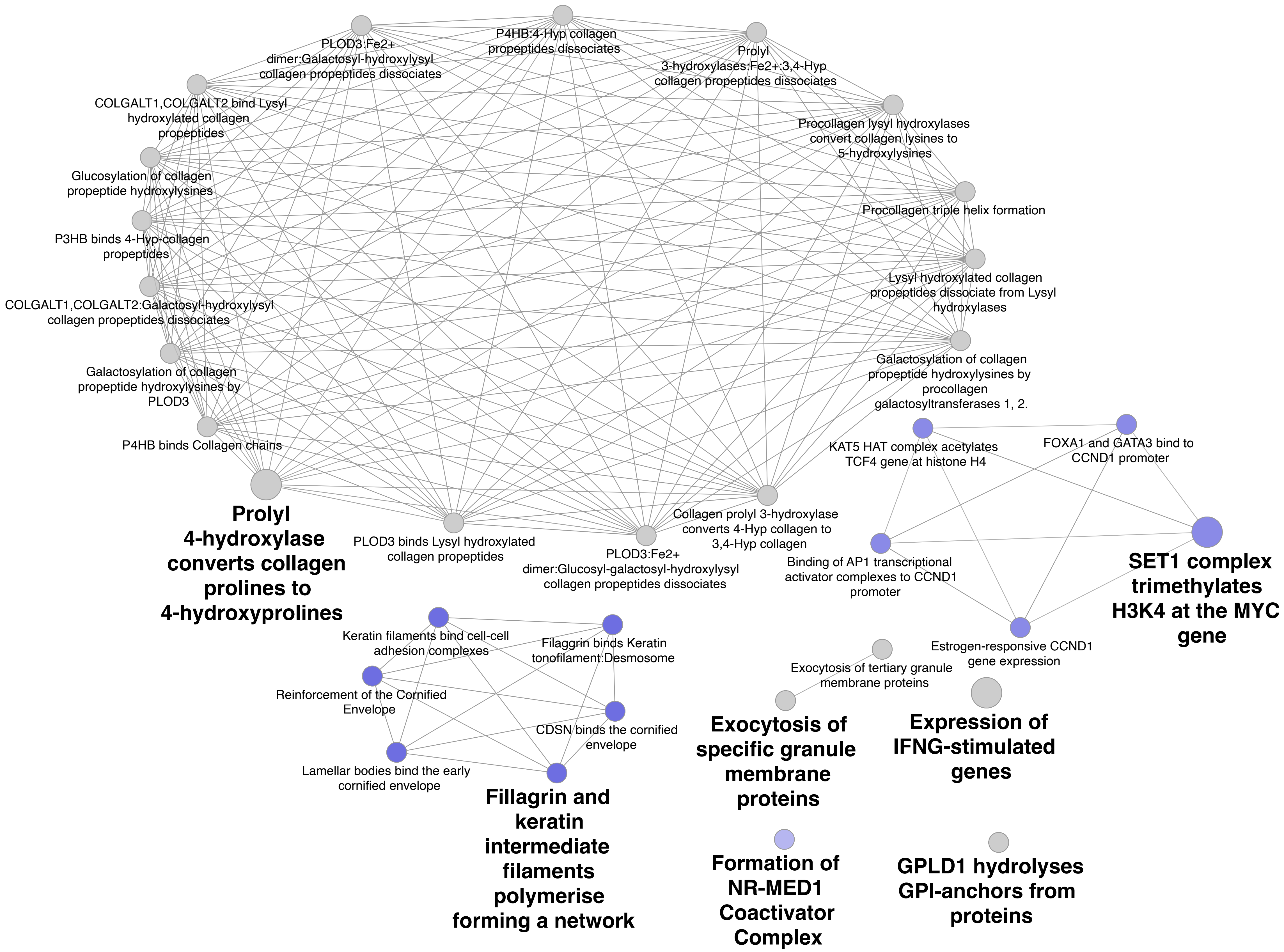


**Supplementary Figure 10: ClueGO Reactome reactions network map.** Grey nodes indicate shared terms, blue nodes indicated terms enriched in PP comparison, and green nodes indicate terms enriched in MM comparison.

A

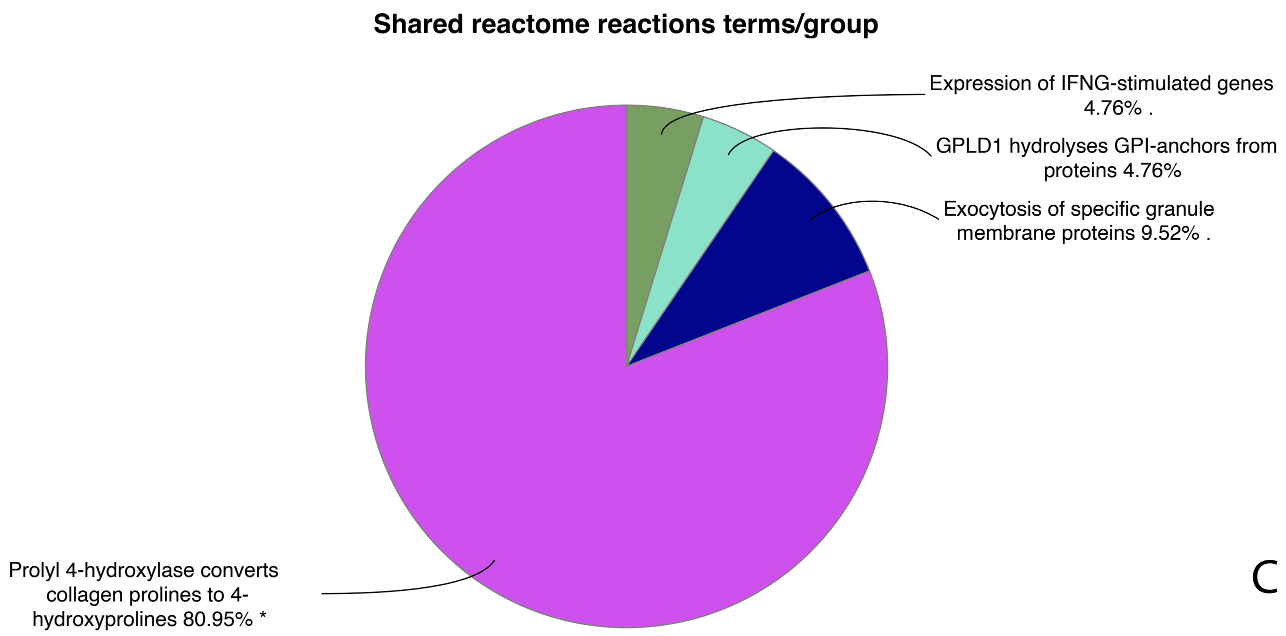


C

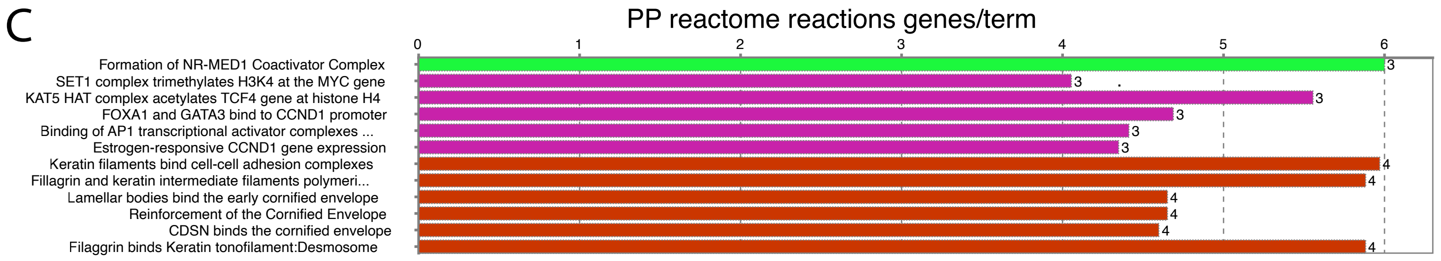


B

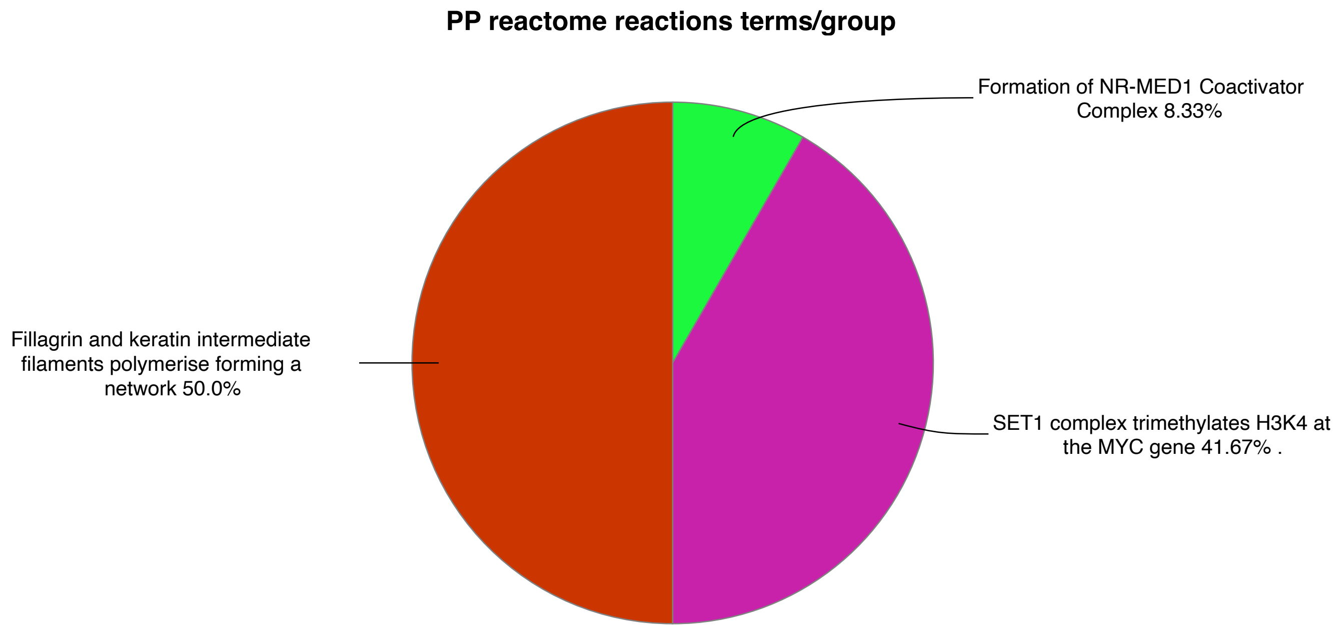


**Supplementary Figure 11: ClueGO Reactome reactions terms and groups.** Pie charts show the proportion of the terms associated with the groups for Shared (A) and PP (B). Histograms indicate number of genes found per term for the PP (C) comparison. No terms were enriched in the MM comparison.

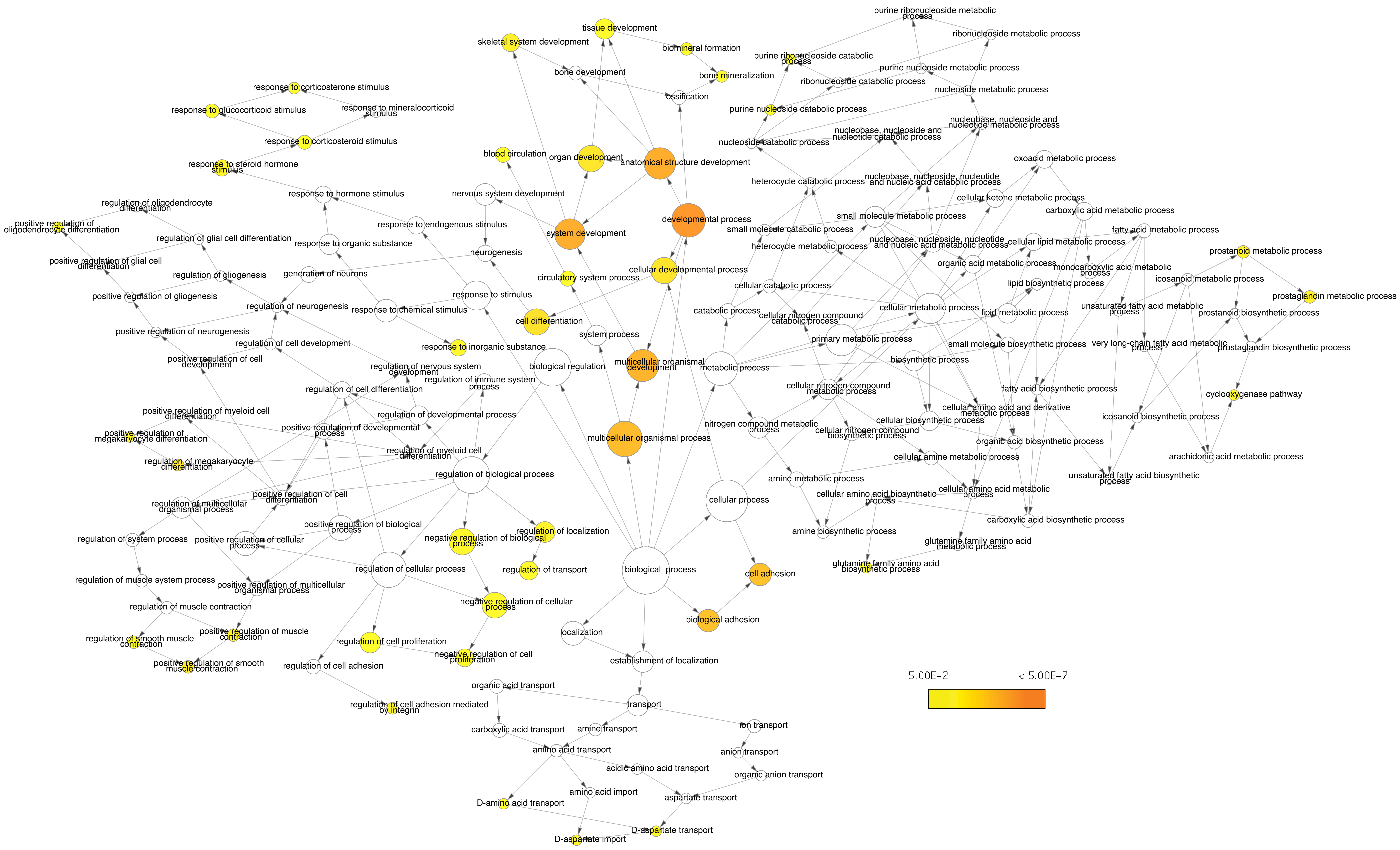


**Supplementary Figure 12: BiNGO network map of Biological process for PP comparison.** Node color indicates the term P value. White nodes are not significant by P value but are included to hierarchically link significant terms.

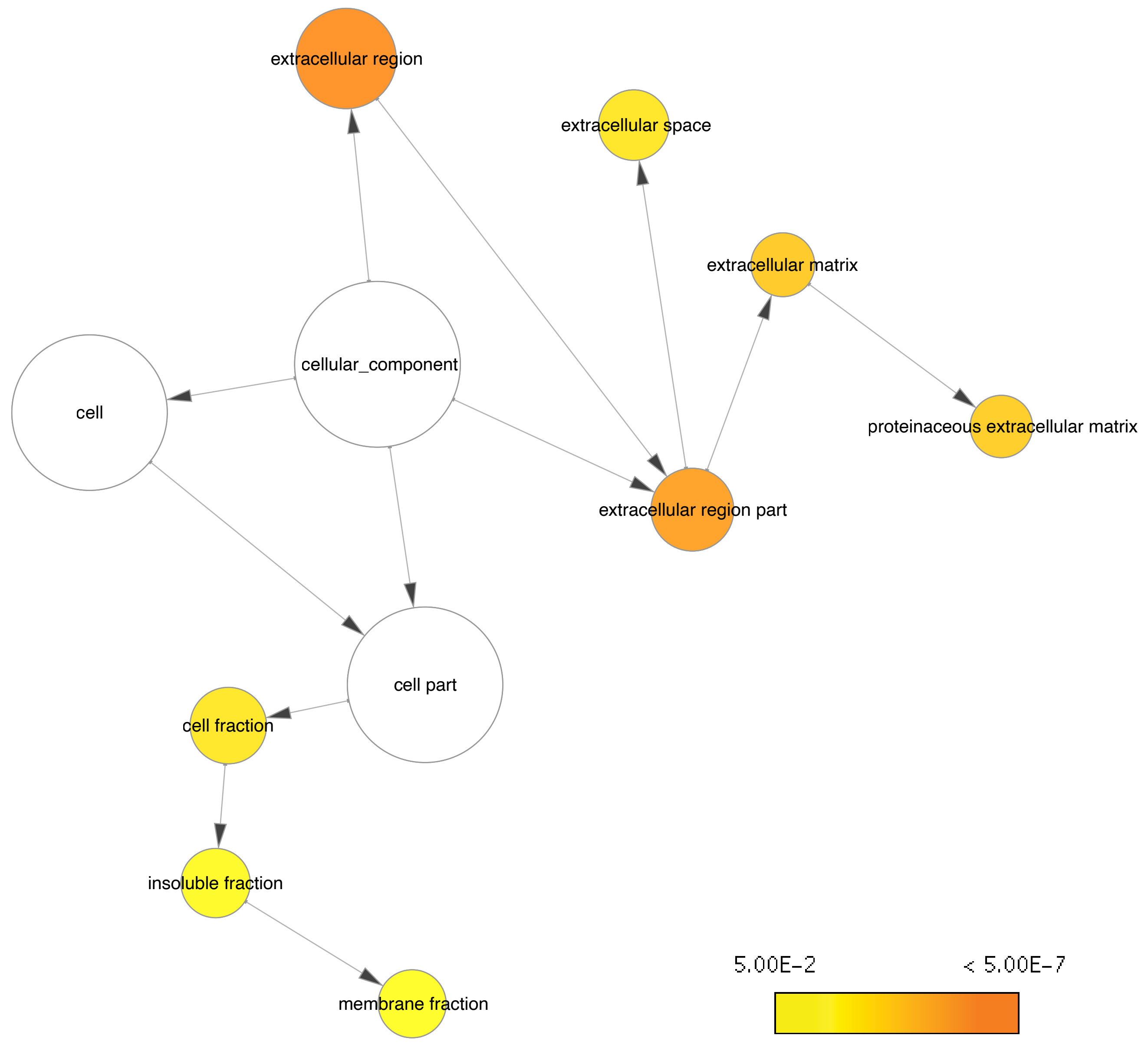


**Supplementary Figure 13: BiNGO network map of Cellular component for PP comparison.** Node color indicates the term P value. White nodes are not significant by P value but are included to hierarchically link significant terms.

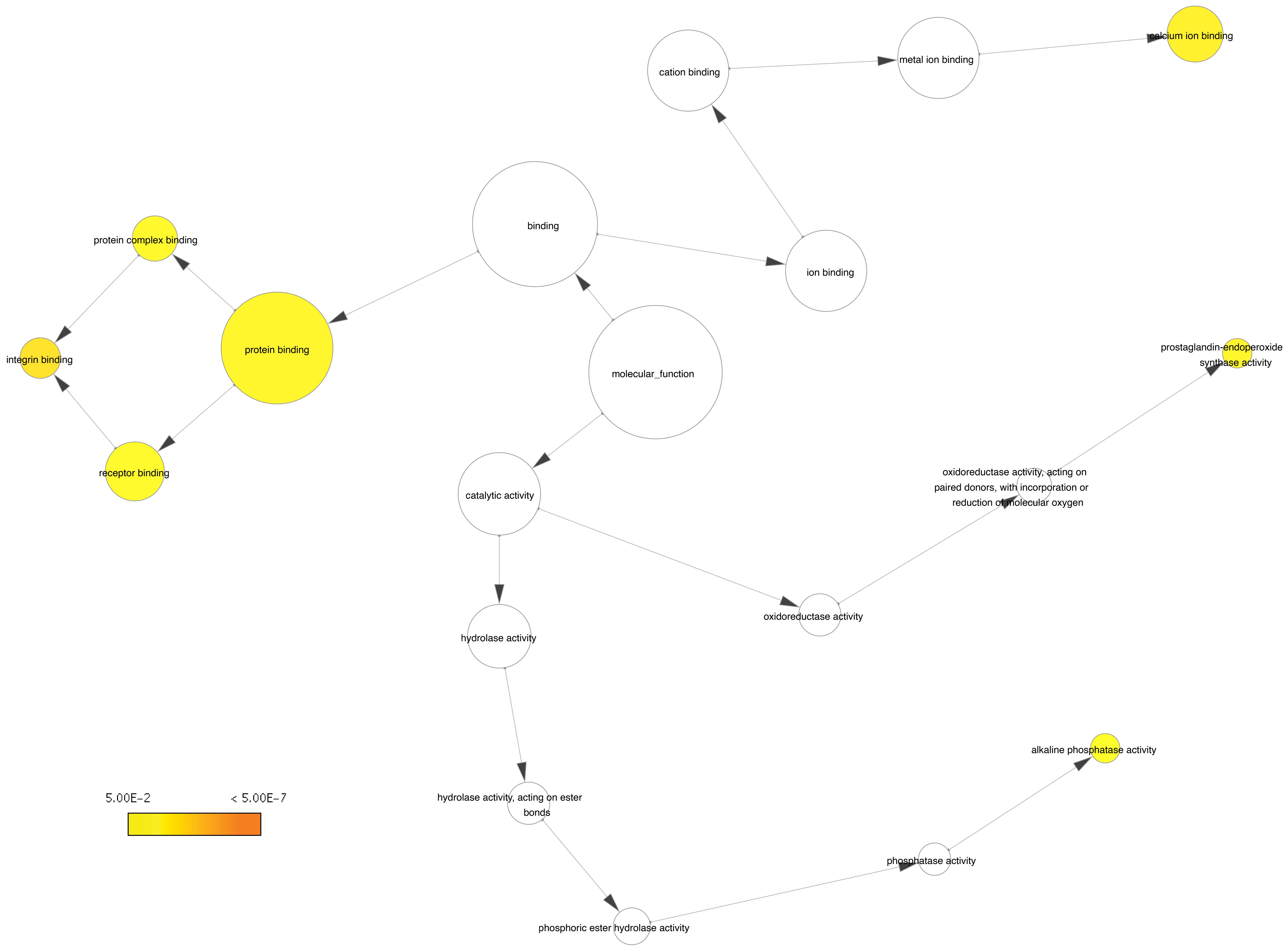


**Supplementary Figure 14: BiNGO network map of Molecular function for PP comparison.** Node color indicates the term P value. White nodes are not significant by P value but are included to hierarchically link significant terms.

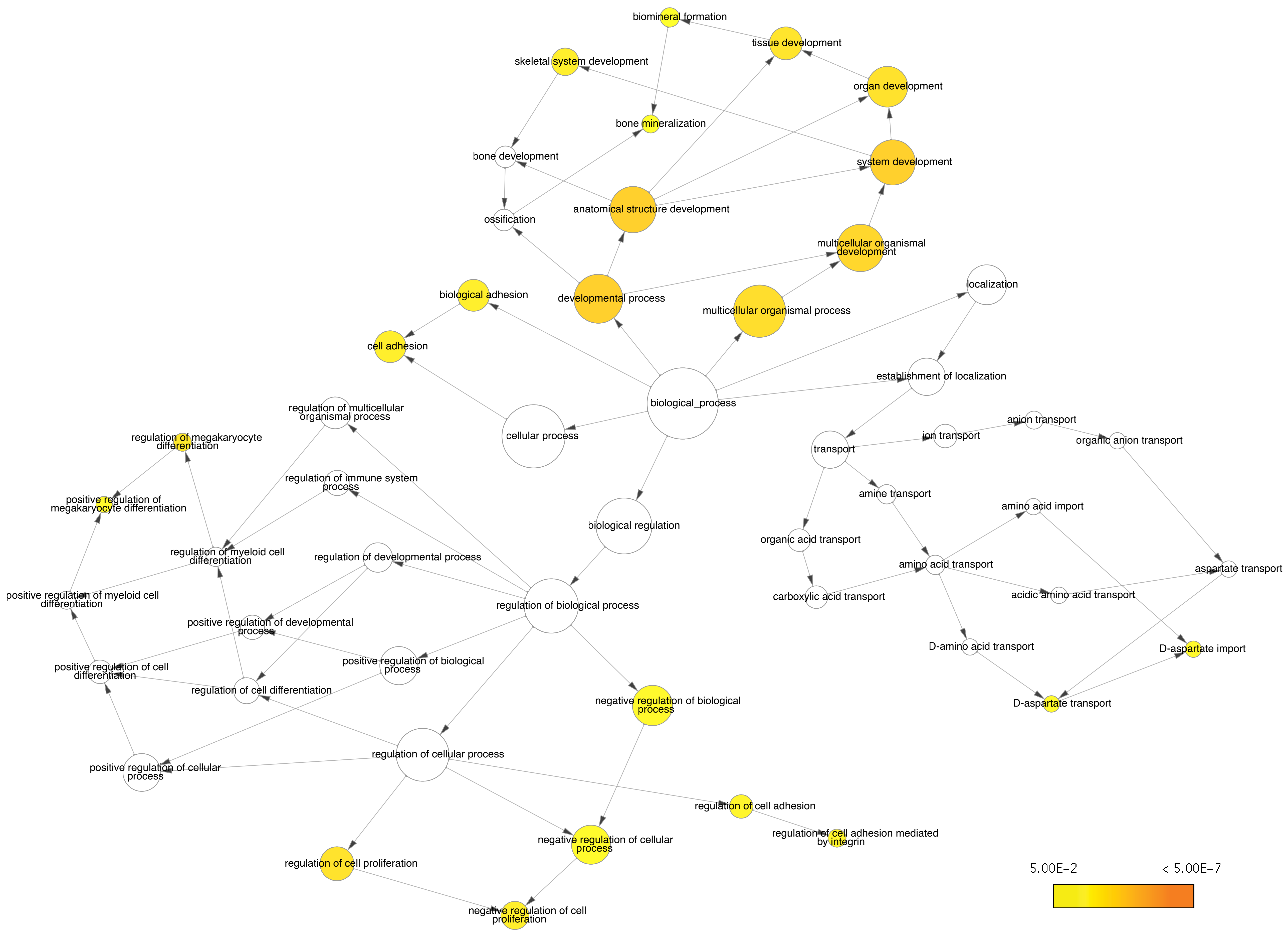


**Supplementary Figure 15: BiNGO network map of Biological process for MM comparison.** Node color indicates the term P value. White nodes are not significant by P value but are included to hierarchically link significant terms.

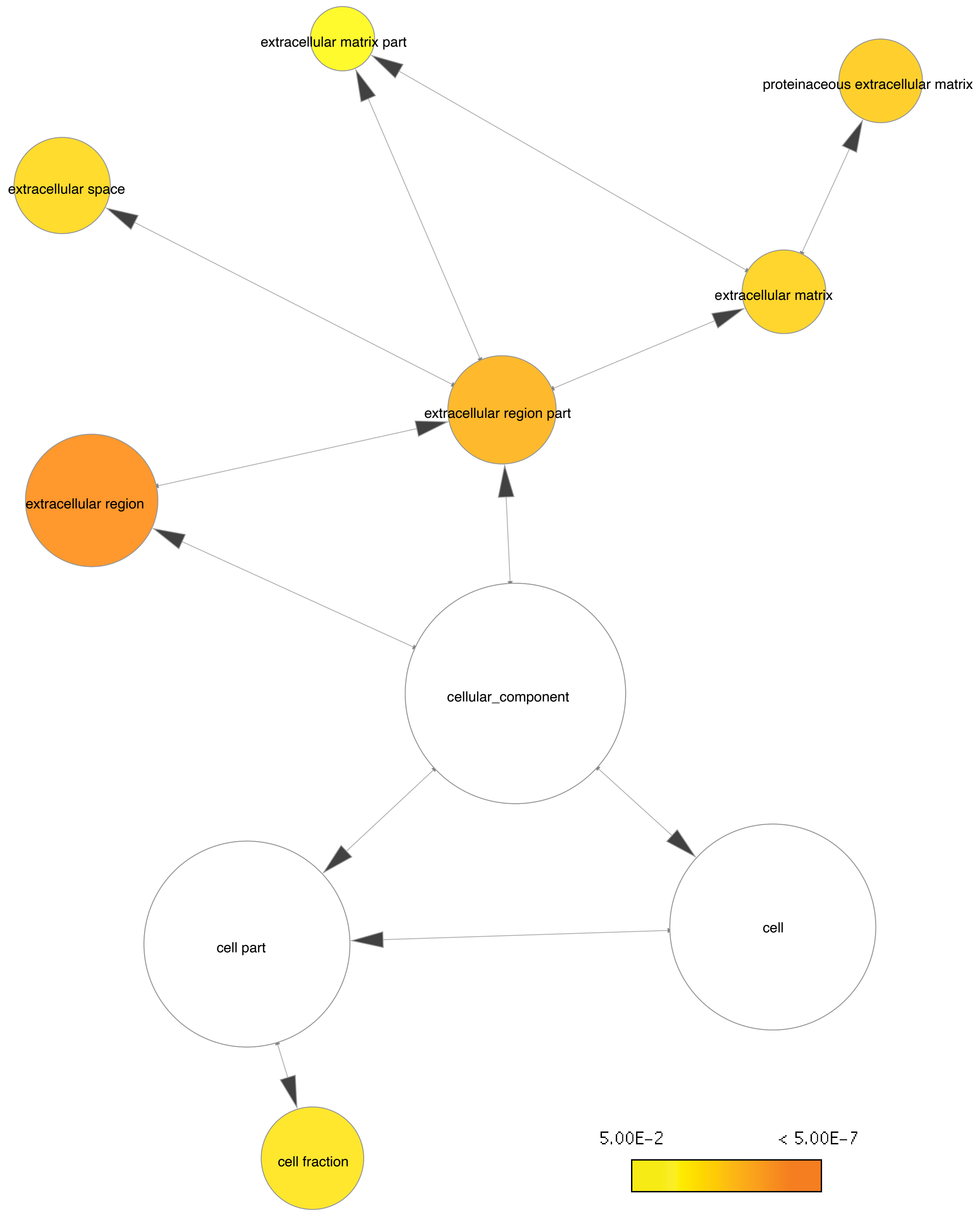


**Supplementary Figure 16: BiNGO network map of Cellular component for MM comparison.** Node color indicates the term P value. White nodes are not significant by P value but are included to hierarchically link significant terms.

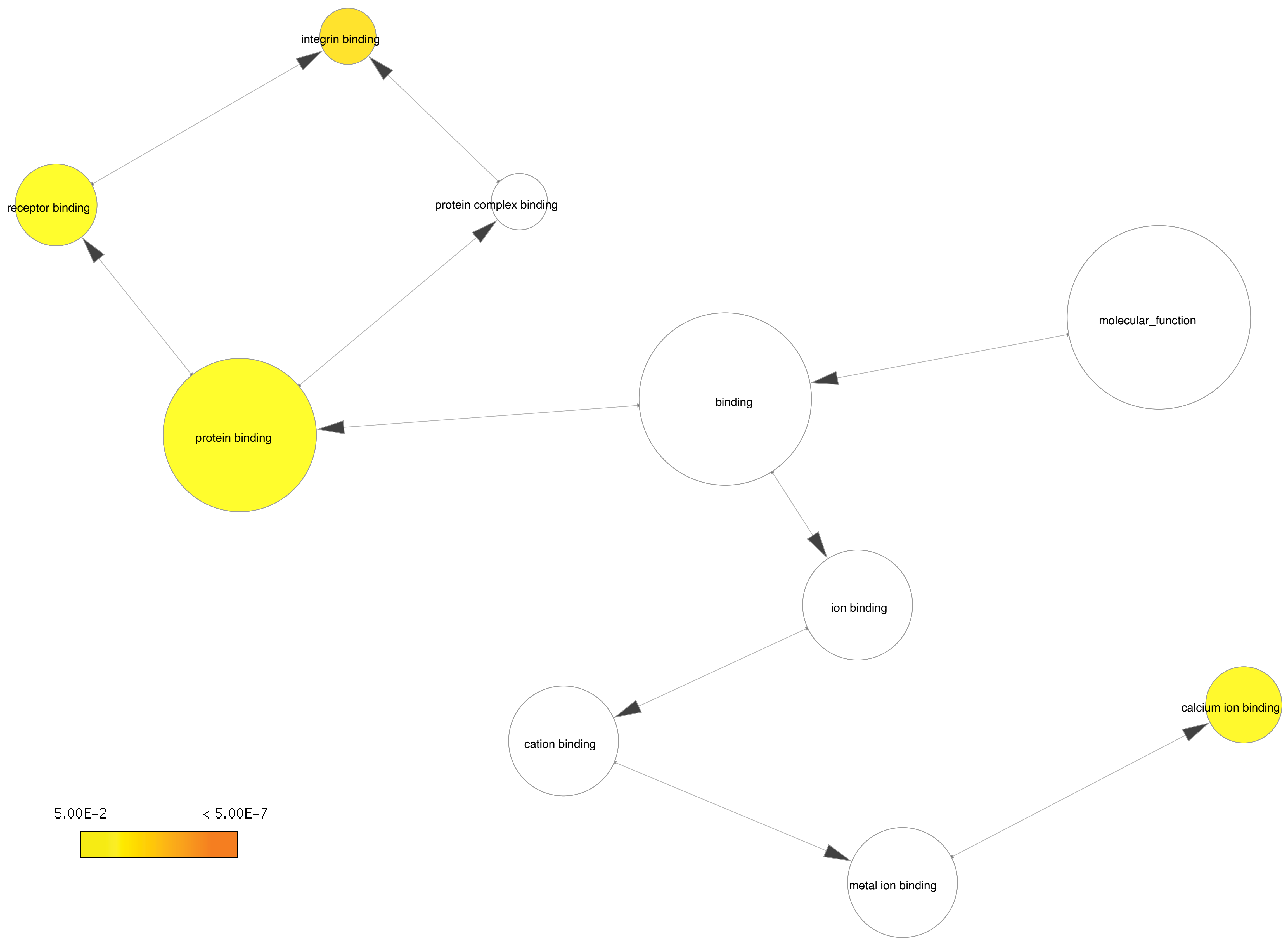


**Supplementary Figure 17: BiNGO network map of Molecular function for MM comparison.** Node color indicates the term P value. White nodes are not significant by P value but are included to hierarchically link significant terms.
